# A novel preclinical secondary pharmacology resource illuminates target-adverse drug reaction associations of marketed drugs

**DOI:** 10.1101/2023.01.31.526400

**Authors:** Jeffrey J Sutherland, Dimitar Yonchev, Alexander Fekete, Laszlo Urban

## Abstract

*In vitro* secondary pharmacology assays are an important tool for predicting clinical adverse drug reactions (ADRs) of investigational drugs. We created the Secondary Pharmacology Database (SPD) by testing 1958 drugs using 200 assays to validate target-ADR associations. Compared to public and subscription resources, 95% of all and 36% of active (AC50 < 1 µM) results were unique to SPD, with bias towards higher activity in public resources. Annotating drugs with free maximal plasma concentrations, we found 684 physiologically relevant novel off-target activities. Furthermore, 64% of putative ADRs linked to target activity in key literature reviews were not statistically significant in SPD. Systematic analysis of all target-ADR pairs identified several novel associations confirmed by publications. Finally, candidate mechanisms for known ADRs are proposed based on SPD off-target activities. Taken together, we present a unique freely-available resource for benchmarking ADR predictions, explaining novel phenotypic activity and investigating clinical properties of marketed drugs.

## INTRODUCTION

Adverse drug reactions (ADRs) are a significant cause of drug discovery and clinical program terminations and post-marketing drug withdrawals^1^. Further, ADRs are a frequent cause of patient drug discontinuation, increasing disease burden for patients and the healthcare system^2^. Anticipating the ADR profile of investigational drugs during lead optimization allows drug discovery teams to pursue strategies for reducing the safety liability while maintaining favorable on-target pharmacological properties.

ADRs mediated by unintended drug activity may involve interaction with one or more targets in the druggable proteome^3^. Despite advances in high-throughput transcriptomic, proteomic or cellular imaging techniques for predicting ADRs^4^, panels of *in vitro* biochemical and cellular assays measuring the effect of drugs on key protein targets retain their pre-eminence in preclinical secondary pharmacology testing^5,6^. However, the number of targets with well-established roles in mediating ADRs is limited. Examples include hERG (KCNH2) for QT prolongation, α1A adrenergic receptor (ADRA1A) modulation for arrhythmia (agonists) or orthostatic hypotension (antagonists), and dopamine D1 (DRD1) antagonism for dyskinesia and tremors^7^. Beyond the hERG channel, lack of scientific consensus on the strength of evidence linking target activity to ADRs may contribute to the high variability in panel composition across pharmaceutical industry^8^.

Prior studies have explored relationships between activity results from biochemical *in vitro* assays and ADRs from marketed drugs^9–12^. These studies have been qualitative in nature (e.g., citing literature implicating the target), were limited to activity results from literature curation available from resources such as ChEMBL^13^ and DrugCentral^14^ and generally used measures of activity potency that did not account for variable human pharmacokinetic properties of drugs, namely the maximal drug exposure (Cmax) at the highest approved dose. Recently, Smit et al^15^ reported the first systematic analysis of safety margin vs. ADR relationships using biochemical activity and human exposure results from ChEMBL, and identified 45 targets with statistically significant relationships vs. human ADRs. Because results from ChEMBL are parsimonious (i.e., most assay vs. compound pairs have no reported results from the literature), the authors used QSAR modelling to fill in missing values and could not account for potential confounding relationships when establishing statistical significance.

Over the course of several years, we have systematically evaluated the activity of 1958 drugs vs. panels of biochemical and cellular *in vitro* assays to create a secondary pharmacology database (SPD). Unusually for such resources, all compounds were tested at 8 or more concentrations, with the concentration resulting in 50% of maximal activity (AC50) available for all tested drug vs. assay pairs. The database reports ca. 150,000 AC50 values for marketed drugs, allowing systematic analysis of target (assay) vs. ADRs reported in databases such as SIDER^16^ and the FDA adverse drug reaction reporting system (FAERS). To our knowledge, the only comparable resource is the Eurofins BioPrint database^17^ which is available by subscription only. Here, we report overall low concordance between results from the SPD (obtained using a limited number of assay protocols for each target) vs. results from ChEMBL and DrugCentral (obtained using a wide variety of such protocols). We illustrate the utility of the database by identifying novel drug activities that may account for drugs’ therapeutic benefit and/or limitation. We used the SPD to identify putative novel target vs. ADR associations via systematic analysis and explain known ADRs via target activities not previously reported in public resources. Beyond the present work, the SPD has broad utility for drug safety and mechanism-of-action investigations, and phenotypic activity deconvolution for novel drug activity in cellular models.

## RESULTS

### *In vitro* safety pharmacology database and comparison to external sources

*In vitro* safety pharmacology assays are used to reveal potential clinical ADRs of low molecular weight compounds during lead optimization^8^. To interpret these results in the context of marketed drug safety profiles, we tested 1958 unique drug substances across 200 safety pharmacology assays, resulting in 147 653 concentration-response curves (Table S1, Table S2). As some of these results were repeated measurements on different dates and/or different lots of drug substance, results were summarized into 121 097 unique drug vs. assay pairs (Dataset S1). The number of assays per drug ranged from 20 to 161, with a median of 66. Thus, SPD represents an unprecedented resource for investigating on- and off-target pharmacology of marketed drugs.

The database was completed over the course of several years. Changes in assay formats (e.g., radioligand binding vs. luminescence), internal vs. external sourcing and other factors resulted in multiple assays for a given target and mode (agonist, antagonist, inhibitor). To simplify data analysis and maximize the number of tested drugs for a given target and mode, assays were merged into 168 assay groups by analyzing concordance of repeated measurements. Assay groups ranged from 1 to 4 assays, and often combine similar assays performed internally vs. contract research organizations (CROs). The number of unique drugs tested per assay group ranged from 30 to 1942, with a median of 794 drugs per assay.

*In vitro* assay results for marketed drugs published in the literature are available in several open-access resources such as ChEMBL^13^ and DrugCentral^14^, and from commercial providers of results curated from journals. Results from our database were cross-referenced to these sources, revealing low overall coverage of drug-target pairs, especially for inactive results (Fig. 1A). Further, quantitative agreement of AC50 values is modest (Fig. 1B-C), possibly because of heterogeneity in methods used to assess pharmacological activity when aggregated across publications. For drug-assay pairs with results in ChEMBL, 66% of SPD AC50 values ≥ 10 µM have a median ChEMBL AC50 < 10 µM. Conversely, 82% of SPD AC50 values < 0.1 µM are reported to be similarly potent in ChEMBL. These observations are consistent with publication bias towards positive or active findings.

**Fig. 1.**
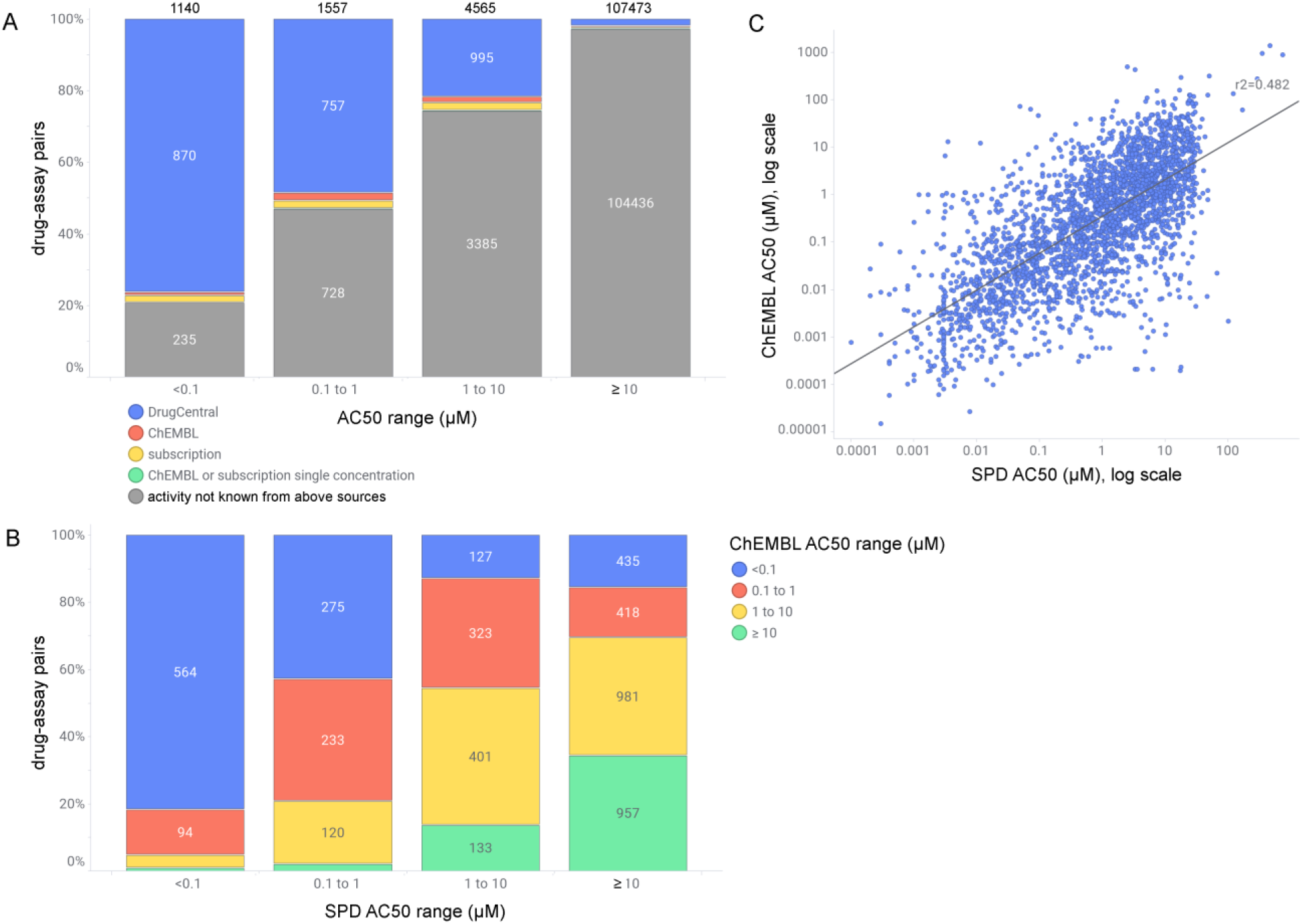
Safety pharmacology results vs. literature-based resources. A) Distribution of AC50 values from SPD by AC50 range, where highly active results are denoted as AC50 < 0.1 µM and inactive results as AC50 ≥ 10 µM. Drug-assay pairs were cross-referenced to resources containing results curated from the biomedical literature: DrugCentral AC50 < 10 µM (blue), ChEMBL AC50 < 10 µM (red), subscription resources AC50 < 10 µM (yellow), or single concentration activity > 50% in either ChEMBL or subscription resources (green). Resources were labelled hierarchically, i.e., activities reported in DrugCentral are mostly available in ChEMBL and other resources. B) qualitative comparison of median ChEMBL vs. SPD AC50 values for 5106 drug-assay pairs; SPD results with AC50 qualifier ‘>’ (AC50 greater than max concentration tested) are shown as ≥ 10 µM; C) quantitative comparison of median ChEMBL vs. SPD AC50 values for 2700 drug-assay pairs where the SPD AC50 qualifier is ‘=’ (i.e., measurable activity); Pearson R^2^ = 0.48.

### Assessing clinical relevance of drug-assay associations

The clinical relevance of results from *in vitro* safety pharmacology panels is commonly assessed by calculating a safety margin, or the ratio of *in vitro* AC50 and the therapeutic free plasma concentration at the highest approved dose^5^. To calculate safety margins from the SPD, we compiled human plasma Cmax and plasma protein binding (PPB) results from several sources, obtaining free Cmax estimates for 937 drug substances (Table S1; Methods). Across all assays, 6783 drug vs. assay safety margins were calculated and distinguished by on-target activities (i.e., the assay measures activity at the drug’s target), off-target activity that is known (in DrugCentral, ChEMBL or subscription resources), and potentially novel off-target activities in the SPD (Fig. 2A). The median on-target safety margin was 2.4, vs 80 for known off-target activities and 353 for novel activities. This suggests that a large proportion of off-target activities from biochemical assays would not manifest in ADRs at clinically relevant exposures.

**Fig. 2.**
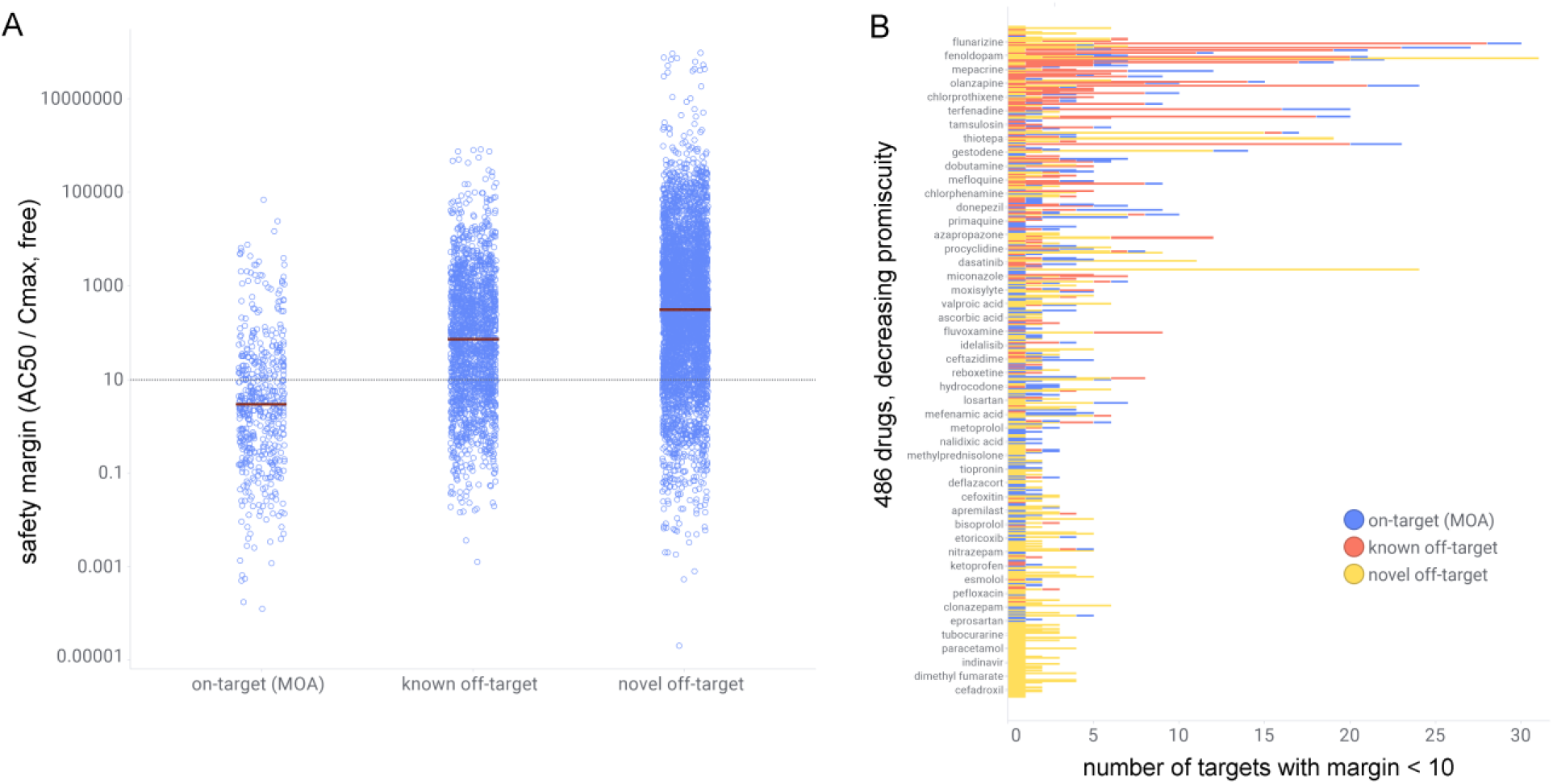
Physiological off-target activities. A) Safety margin (AC50 vs. therapeutic free plasma concentration) of drug-assay pairs by type: on-target activities, off-target activities known in literature-based resources, novel off-target activities. B) Distribution of target activities with safety margin < 10 for 486 drugs, sorted by decreasing overall promiscuity (% of results with AC50 < 10 µM)

According to the free-drug hypothesis, biochemical activity from *in vitro* assays becomes physiologically relevant when the safety margin approaches 1. Overall, 28% (122/429) of on-target margins in our database exceed 10 and 12% (51/429) exceed 50. To investigate target-dependence of safety margins, we tabulated the median value for mechanisms (target and mode; Table S3). Among mechanisms represented by 10 or more drugs, the median margin ranged from 0.2 (SLC6A4, or serotonin transporter) to 5.9 (HTR2A antagonism). Several mechanisms with lower representation had median margin exceeding 10, and 23/47 mechanisms have 25% or more of drugs with margins of 10 or higher.

We labelled as potentially physiological all off-target activities with margin of 10 or less, resulting in 517 known and 684 novel physiological off-target activities. Drugs with higher overall promiscuity, defined as the percentage of assay groups with AC50 ≤ 10 µM, contributed a significant portion of known and novel physiological off-target activities (Fig. 2B). For instance, the promiscuous DRD2/HTR2A antagonist loxapine binds the HTR2B receptor with activity of 6 nM, a result which is not reported in the sources we considered. To identify novel off-target activities with potential impact on disease unrelated to the on-target activity, we focused on a subset with non-overlapping on- vs. off-target indications according to DrugCentral (Table S4). Novel activities that were noteworthy included CNR1 inhibition of losartan (AC50 = 1.2 nM) and glipizide (AC50 = 19 nM), which may contribute to their therapeutic effect in treating hypertension and metabolic syndrome (the CNR1 antagonist rimonabant is used in the management of obesity). Rotigotine, a dopamine agonist used for Parkinson’s disease, was found to be an ADRA1A (α_1a_) agonist (AC50 = 3.3 nM); it has been described as an α_2b_ agonist in a journal not curated within the sources used in our analysis^18^. Clinical studies have shown increased systolic blood pressure in treated patients, consistent with α_1a_ agonism^19,20^. Citalopram, a selective serotonin reuptake inhibitor (SSRI), was found to inhibit the histamine H2 receptor (HRH2; AC50 = 350 nM). Depression and the use of tricyclic antidepressants increase the incidence of gastro-esophageal reflux disease (GERD)^21^. Reports of citalopram use, and GERD are mixed, and clinical studies suggest direct effects (rather than altered pain perception) on esophageal function. HRH2 antagonists (e.g., ranitidine, cimetidine) reduce gastric acid secretion and ameliorate GERD symptoms. Therefore, HRH2 inhibition by citalopram may contribute to its observed effects on the digestive system. Zolpidem, a GABA agonist used for treating insomnia, inhibited CHRM1 (AC50 = 0.21 µM), potentially contributing to its observed effect on dystonia^22^.

### Evaluation of literature-reported target – adverse drug reaction relationships

Variability in the composition of safety pharmacology assay panels suggests that many target vs. clinical ADR associations are not fully understood^8^. Associations reported in the biomedical literature are summarized in three reviews^5,9,23^. We utilized the SPD to evaluate the significance of associations involving 60 targets, each listed as risk factors for 1 to 34 ADRs coded using MedDRA preferred terms. To characterize the strength of such associations, we correlated drug activity measured in our assays vs. clinical ADRs according to SIDER^16^ and FAERS as summarized in DrugCentral^24^. Drug activity was represented as (unadjusted) AC50 values, 2) ratio of AC50 vs. Cmax,tot (i.e., “total” safety margin), and 3) ratio of AC50 vs. Cmax, free (“free” safety margin), and correlated vs. presence or absence of ADRs using the Kruskal-Wallace (KW) test (Fig. S1). Because we performed multiple statistical tests per target-ADR pair, associations were classified as significant (p≤0.001), marginal (0.001 < p ≤ 0.05), not significant (p > 0.05) or not tested (associations with fewer than 10 positive or 50 negative drugs for the ADR). We also imposed a minimal threshold ROC AUC ≥ 0.6 for distinguishing positive vs. negative drugs for a given ADR (Fig. 3A). Across 719 tested associations, 240 (33%) were significant and a further 20 (3%) were marginal. The proportion across targets varies significantly (Fig. 3B). A large percentage of associations were confirmed for adrenergic receptors (e.g., ADRA1A; 15/15), muscarinic receptors (e.g., CHRM1; 30/34), 5-HT receptors (e.g., HTR1A; 20/20 – the notation indicates number significant + marginal / number tested). Our evaluation of target-ADR relationships from Bowes et al^5^, which represents a consensus of safety pharmacology targets across several pharmaceutical companies, is summarized in Table 1, with full results presented in Table S5.

**Fig. 3.**
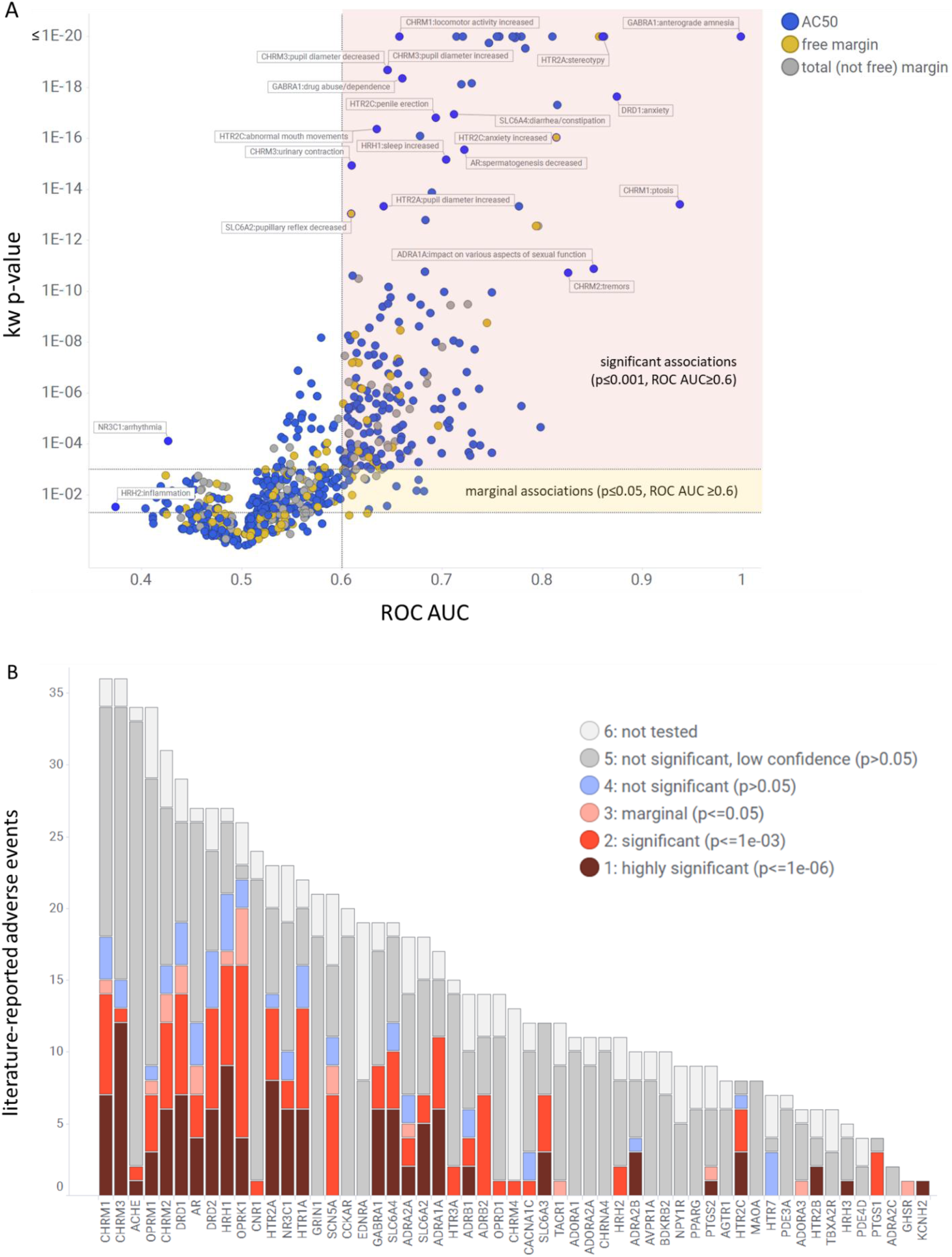
Statistical significance of literature-reported target vs. ADR associations. A) Assessing significance by comparing KW p-value vs. ROC AUC for 884 associations, using the activity measure AC50, free margin or total margin with smallest p-value for each association; regions considered significant (p≤0.001 and ROC AUC≥0.6) or marginal (p≤0.05 and ROC AUC≥0.6) are indicated with shading; a small number of associations having p ≤ 0.05 and ROC AUC < 0.5 denote associations where the direction is opposite to that reported in the literature (drugs with activity have lower probability of exhibiting the ADR); B) Total number of adverse drug reactions reported across targets, distinguished by level of statistical significance observed.

**Table 1.** Investigation of clinical adverse drug reactions attributed to activity in safety pharmacology targets.

| target | mode | % sig. (total)<sup>a</sup> | Significant ADRs<sup>b</sup> |
| --- | --- | --- | --- |
| **ACHE** | inhibition | 6 (33) | salivation ↑, tingling/weakness in limbs |
| **ADRA1A** | activation | 67 (6) | BP ↑, cardiac arrhythmia, pupil diameter ↑, smooth muscle contraction |
| **ADRA1A** | inhibition | 78 (9) | BP ↓, dizziness, heart rate ↑, impact on various aspects of sexual function, orthostatic hypotension, retrograde ejaculation, smooth muscle tone ↓ |
| **ADRA2A** | activation | 44 (9) | heart rate ↓, pupil diameter ↑, sedation |
| **ADRA2A** | inhibition | 20 (5) | heart rate ↑ |
| **ADRA2B** | activation | 43 (7) | heart rate ↑, sedation, skeletal muscle tremor |
| **ADRB1** | activation | 38 (8) | bronchospasm, heart failure, ventricular fibrillation |
| **ADRB1** | inhibition | 50 (2) | heart rate ↓ |
| **ADRB2** | activation | 67 (9) | bronchospasm, cardiac arrest ↑, heart failure, heart rate ↑, QTC interval ↑, skeletal muscle tremor |
| **ADRB2** | inhibition | 50 (2) | BP ↓ |
| **AGTR1** | inhibition | 14 (7) | respiratory distress syndrome |
| **AR** | activation | 43 (7) | androgenicity in females, prostate carcinoma ↑ |
| **AR** | inhibition | 32 (19) | breast carcinoma ↑, insulin resistance, mastodynia, sexual dysfunction, spermatogenesis ↓ |
| **CACNA1C** | activation | 33 (3) | locomotor activity ↓ |
| **CHRM1** | activation | 38 (29) | blurred vision, centrilobular liver congestion, exhaustion, heart rate ↑, irritability, locomotor activity ↓, ptosis, pupil diameter ↓, salivation ↑, tachycardia |
| **CHRM1** | inhibition | 80 (5) | blurred vision, cognitive function ↓, heart rate ↑, locomotor activity ↑ |
| **CHRM2** | activation | 45 (22) | blurred vision, cardiac action potential duration decrease, exhaustion, heart rate ↓, heart rate ↑, PR interval ↓, pupil diameter ↓, salivation ↑ |
| **CHRM2** | inhibition | 80 (5) | cardiac conduction ↓, heart rate ↑, tachycardia, tremors |
| **CHRM3** | activation | 20 (25) | blurred vision, bronchoconstriction, bronchospasm, heart rate ↑, pupil diameter ↓, urinary contraction |
| **CHRM3** | inhibition | 89 (9) | blurred vision, constipation, dry mouth, GI motility ↓, interferes with ocular accommodation, intestinal transit ↓, pupil diameter ↑, salivation ↓↑ |
| **CHRM4** | activation | 100 (1) | pupil diameter ↓ |
| **CNR1** | activation | 6 (17) | drug abuse/dependence↓ |
| **DRD1** | activation | 50 (16) | arousal ↑, drug abuse/dependence, dyskinesia, hypotension, locomotor activity ↓, locomotor activity ↑, psychosis |
| **DRD1** | inhibition | 80 (10) | anxiety, coordination disorders, dyskinesia, locomotor activity ↓, parkinsonian symptoms (tremors), parkinsonism, suicidal intent |
| **DRD2** | activation | 56 (16) | body temperature ↓, drowsiness, drug abuse/dependence, fainting, GI transit ↓, hallucinations, locomotor activity ↓, locomotor activity ↑, stereotypy |
| **DRD2** | inhibition | 50 (8) | drowsiness, GI motility ↑, locomotor activity ↓, orthostatic hypotension |
| **GABRA1** | activation | 53 (15) | anterograde amnesia, ataxia, dizziness, drug abuse/dependence, locomotor activity ↓, memory ↓, sedation, sleep ↑ |
| **GABRA1** | inhibition | 50 (2) | convulsions |
| **HRH1** | activation | 40 (15) | BP ↓, drinking ↑, facial swelling, flushing, sweating, tongue swelling |
| **HRH1** | inhibition | 100 (11) | BP ↓, body weight ↑, cardiac arrhythmia, convulsions ↑, GI transit ↓, heart rate ↑, locomotor activity ↑, QTc interval ↑, sedation, sleep ↑ |
| **HRH2** | activation | 33 (6) | drinking ↑, heart rate ↑ |
| **HRH3** | inhibition | 25 (4) | sedation |
| **HTR1A** | activation | 71 (14) | adrenocorticotrophic hormone ↑, body temperature ↓, growth hormone secretion ↑, locomotor activity ↓, locomotor activity ↑, pupil diameter ↓, pupil diameter ↑, reduced rapid eye movement sleep, sleep ↑, stereotypy |
| **HTR1A** | inhibition | 50 (6) | dizziness, locomotor activity ↑, anxiogenic |
| **HTR2A** | activation | 71 (17) | agitation, drug abuse/dependence, hallucinations, heart rate ↑, hyperreflexia, myoclonus, psychosis, pupil diameter ↑, schizophrenia, serotonin syndrome, smooth muscle contraction, stereotypy |
| **HTR2A** | inhibition | 33 (3) | sleep ↑ |
| **HTR2B** | activation | 33 (3) | cardiac valvulopathy |
| **HTR2B** | inhibition | 100 (1) | GI transit ↓ |
| **HTR2C** | activation | 71 (7) | abnormal mouth movements, anxiety ↑, convulsions ↑, locomotor activity ↓, penile erection |
| **HTR2C** | inhibition | 100 (1) | drug abuse/dependence |
| **HTR3A** | inhibition | 33 (6) | constipation, GI transit ↓ |
| **KCNH2** | inhibition | 100 (1) | prolongation of QT interval of ECG |
| **NR3C1** | activation | 67 (12) | blood glucose ↑, body weight ↑, glaucoma, hyperglycemia, insulin resistance, muscle mass ↓, osteoporosis, wound repair ↓ |
| **OPRD1** | inhibition | 33 (3) | pain ↑ |
| **OPRK1** | activation | 85 (20) | anxiety ↑, confusion, dizziness, drinking ↑, drug abuse/dependence, dysphoria, eating ↑, GI motility ↓, GI transit ↓, hallucinations, heart rate ↓, heart rate ↑, locomotor activity ↓, sedation, tachycardia |
| **OPRK1** | inhibition | 100 (3) | convulsions ↑ |
| **OPRM1** | activation | 40 (20) | drug abuse/dependence, GI motility ↓, GI transit ↓, pupil diameter ↓, pupil diameter ↑, respiratory depression, sedation |
| **PTGS1** | inhibition | 75 (4) | dyspepsia, gastric bleeding, renal dysfunction |
| **PTGS2** | inhibition | 33 (6) | urinary sodium excretion ↓ |
| **SCN5A** | activation | 60 (5) | cardiac arrhythmia, heart rate ↑, locomotor activity ↓ |
| **SCN5A** | inhibition | 55 (11) | cardiac arrhythmia, GI transit ↓, heart rate ↓, heart rate ↑ |
| **SLC6A2** | inhibition | 47 (15) | constipation, drug abuse/dependence, locomotor activity ↓, locomotor activity ↑, pupillary reflex ↓, QTC interval ↑, urinary hesitancy |
| **SLC6A3** | activation | 50 (2) | coordination ↓ |
| **SLC6A3** | inhibition | 60 (10) | dyskinesia, dystonia, locomotor activity ↑, parkinsonism, psychostimulation, stereotypy |
| **SLC6A4** | inhibition | 59 (17) | anxiety ↑, diarrhea/constipation, dizziness, GI motility ↑, locomotor activity ↓, locomotor activity ↑, sexual dysfunction, sleep ↓, tremor, upper GI transit ↓ |

<sup>a</sup> percent of associations having p-KW < 0.001 (number of associations tested); <sup>b</sup> BP: blood pressure, GI: gastrointestinal; ↑: increased; ↓: decreased

Several targets had no statistically significant associations, despite many potential ADRs reported in the literature. Lack of significance might be due to characteristics of the assay data (i.e., few actives, low statistical power), few drugs causing an ADR, biases towards certain ADR types, etc. To investigate further, associations were labelled as “significant” (p < 0.001 and ROC AUC ≥ 0.6) or “non-significant” (all others). We created a Lasso-penalized logistic regression model of these outcomes using several properties, including measures denoting proportion of drugs active in the assay and MedDRA system organ class (SOC) of the ADRs. A small number of variables, including decreasing 5^th^ percentile of AC50 values, increasing count of drugs with AC50 < 1 µM, and ADRs belonging to the SOCs “Nervous system disorders” or “Psychiatric disorders” all increased the probability of a target-ADR association being significant (Fig. S2). These results are intuitive: targets having many drugs with potent AC50 values (the percentile measure), or sub-micromolar actives, and CNS-related ADRs observed when modulating promiscuous GPCRs^5^ are more likely to have significant ADRs.

The model class assignments (“likely significant” or “likely non-significant”) can be viewed as a prior likelihood of validating a target-ADR association, given characteristics of the assay data and the ADR class. When evaluating the predictions for 459 literature associations having p > 0.05 or ROC AUC < 0.6, 414 (90%) were assigned the “likely non-significant” class. These are literature-reported associations identified by the model as having low likelihood of being significant, given the dataset. Conversely, 45 associations with p > 0.05 were identified by the model as “likely significant” (i.e., dataset characteristics should enable validation of a true association). These include both serious ADRs (heart failure for ADRA2B, DRD1 and DRD2 activation) and lower severity effects (sleep or memory impairments for several targets).

Safety margins, either based on free^5^ or total Cmax^25^, have been proposed as superior to unadjusted AC50 for predicting ADRs. A practical limitation is that estimated human Cmax is often not available in early lead optimization. The proportion of significant associations was highest when using AC50 as activity measure, more notably when using SIDER as the source of ADR annotations. (Fig. S3A). This remained true when using the ROC AUC rather than p-value for selecting significant associations, a measure which should be less sensitive to the smaller sample sizes for margin-based activity measures (Fig. S3B). Even for target-ADR pairs significant on both AC50 and free margin, the strength of association is generally higher for AC50-based activity measures (Fig. S4).

### Systematic evaluation of target vs. clinical adverse drug reaction relationships

The 743 literature-derived target vs. ADR associations represent a small proportion of all possible such associations. We systematically evaluated all possible target vs clinical ADR annotations from SIDER and FAERS. To increase power to detect associations for less common ADRs, we also modelled assay vs. MedDRA high terms (HT) and group terms (HG) that combine related preferred term (PT) ADRs. In total, we examined 562 744 relationships: activity measures 1) (unadjusted) AC50 values in µM, 2) total margin and 3) free margin for each of 124 assays and 2647 MedDRA terms with sufficient representation in SPD. Overall, 1992 assay vs. MedDRA pairs met the statistical criteria of ROC AUC ≥ 0.7 and KW p-value < 1e-06 on one or more activity measures (Table S6). This included 671 associations using HT or HG terms, of which 560 had one or more significant children (e.g., HG vs. HT or HT vs. PT) terms. Approximately 70% had higher ROC AUC and/or smaller p-value compared to the most significant child term. To limit redundancy, we focused further analysis on 1321 PT and 111 non-redundant HT/HG-based associations.

Only 26% of the 1432 assay vs. ADR associations overlapped with those reported in the literature, as described above, suggesting that some might be novel (Fig. 4A). Mirroring our findings for literature associations, a large proportion were significant only on AC50, compared to margin-based activity measures (Fig. 4B). However, we noted that the proportion supported by a margin-based measure increased with the proportion of active drugs that were on-target, suggesting that margin-based associations were more likely to be plausible (Fig. 4C).

**Fig. 4.**
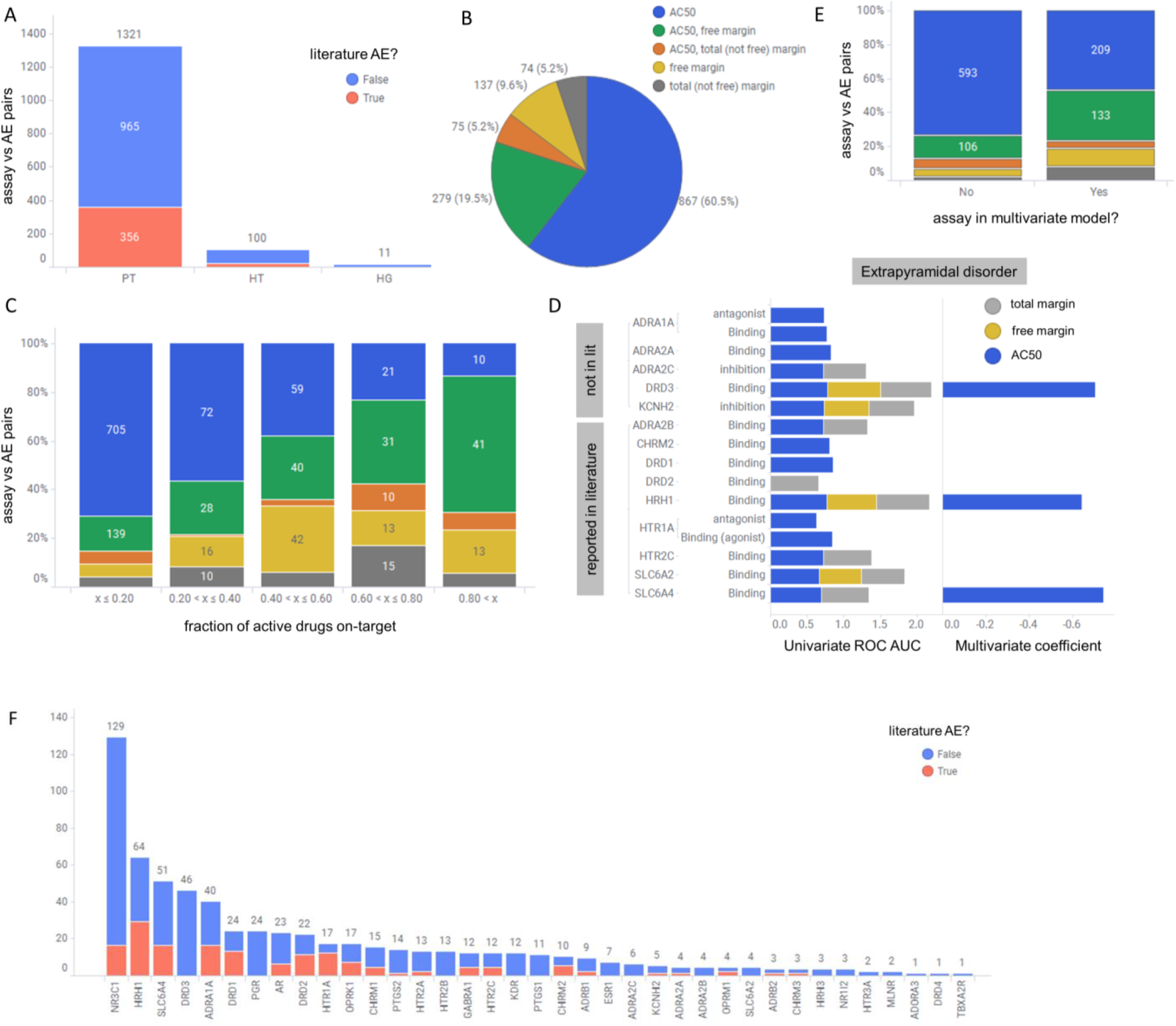
Systematic identification of assay vs. ADR relationships. A) Number of assay vs. ADR pairs having KW p-value ≤ 1e-06 and ROC AUC ≥ 0.7 in either SIDER or FAERS, by MedDRA type (PT – preferred term, HT – high term, HG – group term), distinguished by their presence or absence among the literature reported associations; B) Proportion of associations considered significant by 3 activity measures considered; associations significant by free margin alone, or both free/total margin, are labelled as “free margin”; associations significant by total margin only are labelled as “total (not free) margin”; C) Distribution of associations by activity measure vs. fraction of drugs active in the assay that are “on-target” (i.e., activity at the drugs’ known targets), D) identification of non-redundant assays linked to “extrapyramidal disorder”. The left panel indicates univariate ROC AUC for each assay, showing cumulatively in “stacked” form significant associations for AC50 (blue), free margin (yellow), total margin (gray). The right panel indicates the coefficient in the penalized logistic regression model, i.e., only 3 assay AC50s have non-zero values and are therefore considered non-redundant in explaining ADR risk. E) Comparing assays with non-zero vs. zero coefficients in the logistic model by proportion of significant activity measures; F) Distribution of 631 non-redundant assay vs. ADR pairs by target, distinguished by their presence or absence among the literature associations.

Drugs often modulate off-targets with high sequence similarity (and hence similar binding pockets) vs. the intended target. Statistically significant off-target vs. ADR relationships from univariate analysis may be confounded by the corresponding on-target relationship. For example, the most significant association involving glucocorticoid receptor (NR3C1) binding vs. “Adrenal cortical hyperfunctions” (MedDRA 10001341; p-value = 2e-42, ROC AUC = 0.87) is mirrored by a similar relationship for progesterone receptor (PGR) agonism (p-value = 1e-17; ROC AUC = 0.87). Among 12 drugs causing this ADR according to SIDER and tested in both assays, 8 drugs modulate both NR3C1 and PGR with AC50 < 10 µM; 7 drugs are glucocorticoid antagonists and 1 drug is a PGR agonist. Achieving selectivity is difficult^26^, and whether one or both targets contribute to occurrence of this ADR cannot be determined via univariate statistical analysis.

To distinguish overlapping vs. orthogonal assay contributions to clinical ADR risk, we performed multivariate logistic regression to model the probability of observing a given ADR using the subset of significant assays identified above. Because “novel” high significance assays and known lower significance assays might explain the same variation in ADR risk, we included assays for literature targets that reached the lower significance threshold p ≤ 0.001 (i.e., “significant” per Table S5 but not reaching the combined p ≤ 1e-06 and ROC AUC ≥ 0.7 for inclusion in Table S6). The analysis workflow employing the L1 (or “lasso”) penalty seeks to select the smallest number of variables (assays) resulting in a model with predictive accuracy within one standard error of the best model (with any number of variables). Accordingly, assays retained in the sparse model are likely to represent distinct contributions.

To illustrate, two datasets were modelled to identify non-redundant assays for predicting “Adrenal cortical hyperfunctions”: one from SIDER consisting of 487 drugs annotated as positive (18) vs negative (469) for the ADR, and a second from FAERS consisting of 605 drugs (17 positives vs 588 negatives). These datasets are not identical owing to differences in SIDER and FAERS, with 11 shared positive drugs in both datasets. Measures of AC50, total margin and free margin for NR3C1 and PGR were employed as predictors (3 x 2 variables). The optimal (sparse) model for each dataset was reduced to a single variable, namely the NR3C1 AC50. This suggests that the total and free NR3C1 margins and the PGR endpoints explain the same variation in the ADR risk as the NR3C1 AC50. Put differently, there is no evidence of utility beyond the NR3C1 AC50 to predict this ADR. For “extrapyramidal disorder” (MedDRA 10015832) SIDER annotations, this approach reduced the number of predictive assay + activity measure pairs from 28 to 3, namely SLC6A4, HRH1 and DRD3 binding AC50 values (Fig. 4D); DRD3 is not reported as a risk factor for this ADR according to the literature reviews. Overall, this approach eliminated 801 assay vs. ADR pairs identified as significant by univariate analysis but redundant with other retained (more predictive) assays. Comparing retained vs. eliminated assays shows enrichment towards margin-based measures (Fig. 4E).

In summary, we identified 189 ADRs with a single predictive assay (p-value ≤ 1e-06 and ROC AUC ≥ 0.7; no multivariate modelling) and a further 442 assay vs. ADR pairs with univariate significance and retention in the sparse models. The glucocorticoid receptor (NR3C1), histamine H1 receptor (HRH1), serotonin transporter (SLC6A4), dopamine D3 (DRD3), adrenergic alpha-1A (ADRA1A) and dopamine D1 (DRD1) receptors accounted for the largest proportion of ADRs (Fig. 4F).

### Investigation of potentially novel target vs. ADR relationships

The analytical workflow described above systematically evaluated all possible assay (target) vs. ADR relationships, identifying 631 associations that met stringent statistical criteria (p-value ≤ 1e-06 and ROC AUC ≥ 0.7) and were not redundant with known risk factors (i.e., retained via the sparse modelling). As validation of this approach, 149 associations (24%) were from the literature reviews described above. This suggests that a subset of the remaining 482 associations may represent novel target vs. ADR risk factors.

Amongst these, the glucocorticoid receptor (NR3C1) accounted for many associations. Even though these were not all listed in the literature reviews, immune suppression that results from activity at NR3C1 is well recognized, and most of these interactions are on-target. Activity linked to the serotonin transporter (SLC6A4) similarly shows a range of ADRs associated with the indication of mood disorders that are treated with SSRI drugs.

We searched the literature for associations having at least 20% of drugs active against vs. off-target and that seemed plausible to us (Table 2). For example, ADRA2C inhibition was associated with auditory hallucination and paranoia in FAERS, on both AC50 and margin measures. A mouse knockout model showed increased startle reflex and aggression^27^. Differential gene expression^28^ and genetic alterations^29^ in ADRA2C also suggest a role in schizophrenia. DRD2 and DRD3 binding were associated with mammary and menstruation-related ADRs in SIDER^30–32^. PGR agonism was associated with hyperpigmentation disorders in SIDER, with drugs acting both on-target (medroxyprogesterone, progesterone) and off-target (NR3C1 modulators). A recent report describes the effect of asoprisnil, a selective PGR modulator, on melanocytes^33^.

**Table 2.** Selected assays vs adverse drug reaction relationships from univariate and multivariate analyses with support in the biomedical literature.

| **assay name** | **MedDRA name** | **KW p-value** | **ROC AUC** | **significant measures** | **significant sources** | **Literature<sup>a</sup>** |
| --- | --- | --- | --- | --- | --- | --- |
| ADRA1A Binding (NIBR assay) | Eosinophilia | 3.6E-08 | 0.85 | total (not free) margin | FAERS | 32194050 |
|  | Tardive dyskinesia | 1.9E-23 | 0.87 | AC50, free margin | SIDER | 39,40 |
| ADRA2C inhibition | Hallucination, auditory | 1.1E-11 | 0.86 | AC50, total (not free) margin | FAERS | 27–29 |
|  | Paranoia | 1.2E-09 | 0.82 | AC50, free margin | FAERS | 27–29 |
| AR Binding (agonist) | Erythema nodosum | 2.6E-10 | 0.70 | AC50 | SIDER | 27075133 |
|  | Ovarian and fallopian tube cysts and neoplasms | 2.4E-09 | 0.70 | AC50 | SIDER | 30791431 |
|  | Retinal embolism and thrombosis | 6.5E-07 | 0.74 | AC50 | SIDER | 29040227 |
|  | Ovarian neoplasms malignant (excluding germ cell) | 5.9E-12 | 0.82 | AC50 | SIDER | 30791431 |
| CHRM1 inhibition | Micturition urgency | 4.0E-09 | 0.73 | AC50 | SIDER | 35117285 |
|  | Dental caries | 3.9E-07 | 0.73 | AC50 | SIDER | 31289718<br>12974516 |
| CHRM2 Binding | Ileus paralytic | 2.7E-10 | 0.76 | AC50, free margin | SIDER | 14607264 |
| DRD2 Binding | Breast enlargement | 2.4E-08 | 0.72 | AC50 | SIDER | 30–32 |
|  | Menstruation irregular | 2.4E-08 | 0.71 | AC50 | SIDER | 32 |
|  | Salivary hypersecretion | 7.9E-11 | 0.71 | AC50 | SIDER | 16421461 |
| DRD3 Binding | Tardive dyskinesia | 4.3E-31 | 0.90 | AC50, free margin | FAERS, SIDER | 34–36 |
|  | Extrapyramidal disorder | 5.1E-33 | 0.87 | AC50, free margin | FAERS, SIDER | 41 |
|  | Gambling disorder | 2.8E-09 | 0.83 | AC50 | SIDER | 26192187 |
|  | Hyperprolactinemia | 1.6E-17 | 0.86 | AC50, free margin | SIDER | 16169407<br>33854317 |
|  | Blood prolactin increased | 4.2E-14 | 0.88 | AC50, free margin | SIDER | 33854317 |
| HRH1 Binding | Hypomania | 5.0E-17 | 0.88 | AC50, free margin | SIDER | 34572558 |
| HTR2B antagonist | Lethargy | 7.4E-07 | 0.71 | AC50 | SIDER | 30666218 |
|  | Sleep disorders NEC | 1.9E-08 | 0.72 | AC50 | SIDER | 30666218 |
| OPRK1 Binding | Salivary hypersecretion | 2.8E-08 | 0.74 | AC50 | SIDER | Entrez Gene |
| PGR agonist | Hyperpigmentation disorders | 7.8E-13 | 0.71 | free margin | SIDER | 33 |
| SLC6A4 Binding | Generalized tonic-clonic seizure | 1.5E-08 | 0.76 | free margin | FAERS | 31849820 |
|  | Galactorrhea | 2.0E-21 | 0.76 | AC50, free margin | SIDER | 14997175 |
|  | Blood prolactin increased | 2.5E-07 | 0.78 | AC50, free margin | SIDER | 14997175 |

<sup>a</sup> References not discussed in the text are provided as PubMed IDs

More challenging is the interpretation of motor dysfunction ADRs associated with modulation of DRD3 and adrenergic receptors, given the preponderance of evidence implicating DRD1 and DRD2 (Fig. 4D). Even though sparse model building selected DRD3 over DRD2, both targets may explain the same variation in ADR risk. DRD3 gene polymorphisms have been associated with tardive dyskinesia (TD)^34–36^, and DRD3 knockout animals have slightly altered locomotor activity, and enhanced sensitivity to DRD1/DRD2 agonists^37^. Motor dysfunction related ADRs are listed in the FDA label for cariprazine, an atypical antipsychotic selective for the D3 receptor^38^. As such, DRD3 may contribute to the ADR profile of dopaminergic drugs, usually ascribed to DRD2 in the literature. Although TD is strongly linked with abnormalities of the dopaminergic system, there is some evidence from clinical case studies, that 3H-dihydroergocryptine (3H-DHE)-alpha2 adrenergic receptor binding and cerebrospinal fluid norepinephrine (NE) were positively correlated with the severity of TD^39,40^. It remains elusive to link drug-induced TD to engagement with adrenergic receptors with antipsychotic drugs, as most of them have high pharmacological promiscuity with prominent effects at dopamine, histamine and serotonin receptors, also associated with movement disorder side effects. However, it is plausible that combined effects at these receptors include the adrenergic component.

### SPD assay results reveal putative cause of known drug adverse drug reactions

Many drug vs. assay results from the SPD are not described in the pharmacology resources we interrogated. To determine if these novel activities explain known ADRs of marketed drugs, we tabulated 325 drug-target-ADR triples for target-ADR relationships reported in the literature and statistically significant in our analysis (Fig. 5; Table S7; Table S8). The most prevalent class of newly explained ADRs were cardiac and respiratory effects. These include 16 drugs active at OPRK1 with association to “cardio-respiratory arrest”, with AC50 values ranging from 0.35 to 4 µM. A variety of ADRs belonging to movement disorders (e.g. extrapyramidal disorder, Parkinsonism, dyskinesia) were associated with several targets, including OPRK1 (9 drugs), CHRM2 (8 drugs), ADRA1A (5 drugs). We investigated selected examples in further detail (Table 3)

**Fig. 5.**
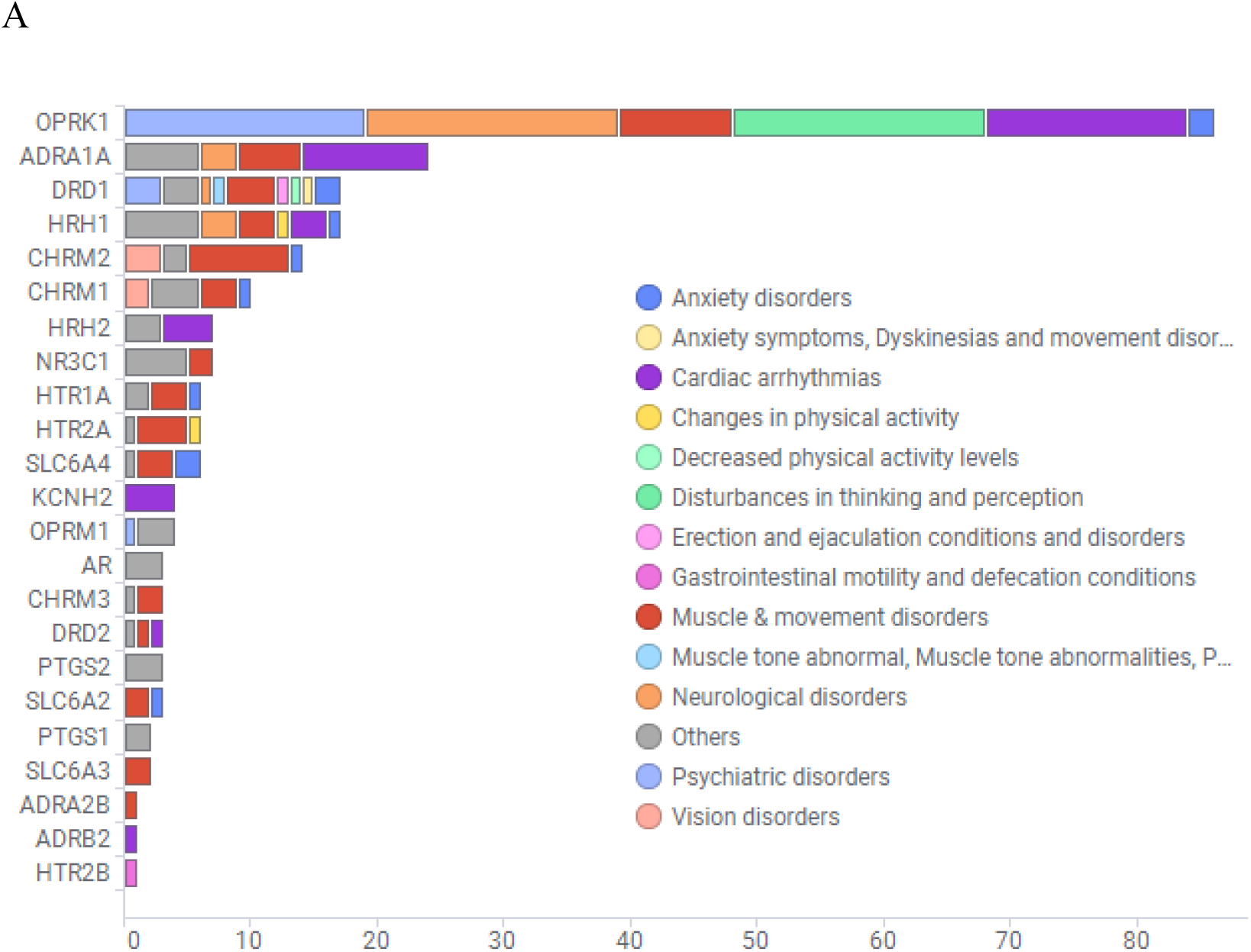
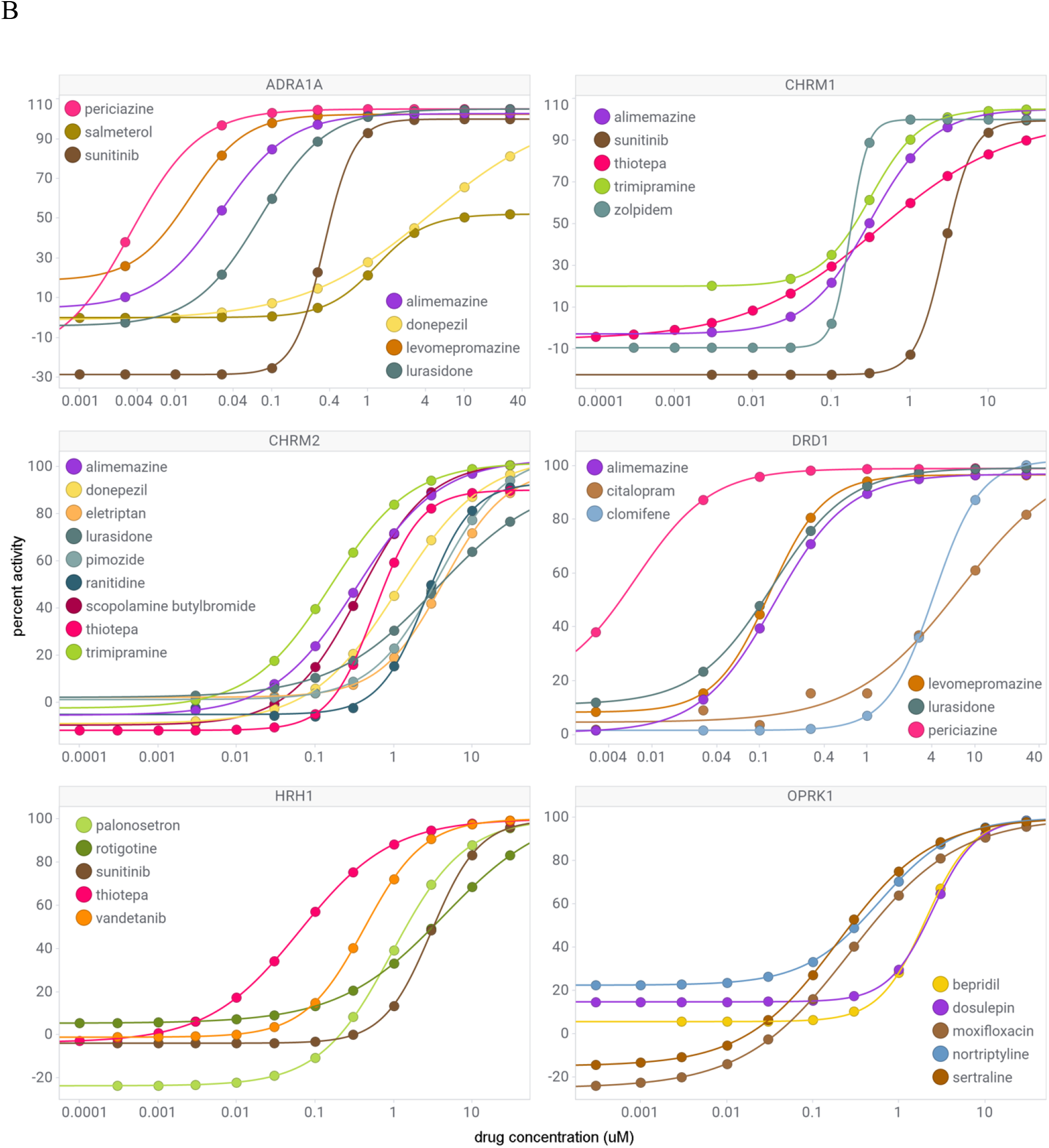
Novel off-target activity as putative mechanisms for known drug vs. ADR associations. A) Number of drug vs. ADR associations with novel off-target activity by target. B) Concentration vs. activity for selected drugs and assays.

**Table 3.** Selected adverse drug reaction newly explained by SPD database.

| **drug (targets)** | **assay target** | **assay AC50 (μM)** | **explained ADRs** |
| --- | --- | --- | --- |
| amiodarone (KCNH2) | HRH1 | 2.4 | akinesia, bradykinesia, decreased activity, disorientation, extrapyramidal disorder, hypokinesia, inappropriate antidiuretic hormone secretion |
| amlodipine (CACNA1C) | ADRB2 | 1.8 | sinus bradycardia |
|  | DRD2 | 2.9 | parkinsonism |
|  | OPRK1 | 1.2 | cardio-respiratory arrest, drug abuse |
| citalopram (SLC6A4) | DRD1 | 3.9 | akathisia, catatonia, choreoathetosis, dyskinesia, ejaculation disorder, extrapyramidal disorder, libido increased, movement disorder, orgasmic disorders and disturbances, parkinsonism, priapism, psychotic disorder, serotonin syndrome, substance related and addictive disorders, tardive dyskinesia, torticollis, trismus |
| citalopram (SLC6A4) | HRH2 | 0.35 | hyperprolactinemia |
| donepezil (ACHE) | ADRA1A | 4.4 | drooling |
|  | CHRM2 | 1.1 | extrapyramidal disorder |
| eletriptan (HTR1B, D, F) | CHRM2 | 4.2 | accommodation disorder |
| fluoxetine (SLC6A4) | OPRK1 | 1.5 | cardio-respiratory arrest, coordination abnormal, delusion, drug abuse |
| lurasidone (DRD2, HTR2A) | ADRA1A | 0.075 | blood prolactin increased, cogwheel rigidity, drooling, muscle rigidity, oculogyric crisis, sexual dysfunction, torticollis |
|  | CHRM2 | 3.5 | dyskinesia |
|  | DRD1 | 0.13 | dyskinesia, sedation, substance related and addictive disorders |
| pregabalin (CACNA2D1,2) | ADRA1A | 3.7 | autonomic nervous system imbalance, blood prolactin increased, dizziness postural, miosis, orgasmic disorders and disturbances, restlessness, sudden death, torticollis, trismus, urinary incontinence |
| quetiapine (DRD2, HTR2A) | OPRK1 | 1.6 | cardio-respiratory arrest, choreiform movements, coordination abnormal, delusion, drug withdrawal syndrome |
|  | OPRM1 | 1.8 | drug dependence, respiratory arrest |
| ranitidine (HRH2) | CHRM2 | 2.6 | dyskinesia |
| sibutramine (SLC6A2,3,4) | HTR2A | 1 | torticollis |
|  | OPRK1 | 4.8 | delusion |
| tramadol (OPRM1) | CHRM2 | 1.5 | dyskinesia, parkinsonism |
| trimipramine (SLC6A2,3,4) | CHRM1 | 0.31 | accommodation disorder, mydriasis |
|  | CHRM2 | 0.16 | accommodation disorder, extrapyramidal disorder, mydriasis, parotid gland enlargement |
| valbenazine<br>(SLC18A2) | HTR1A | 1.4 | tardive dyskinesia |
| zolpidem<br>(GABRA1) | CHRM1 | 0.21 | accommodation disorder, mydriasis |

### Case study: Accommodation disorder and muscarinic activity

Accommodation disorder is one of the most common ADRs associated with muscarinic receptor antagonists^42^. We found several drugs, which do not have known association with muscarinic receptors, but caused accommodation disorder. Zolpidem is a GABA-A receptor agonist with no muscarinic receptor related side effect in its label^44^. However, FAERS data suggest that it is related to accommodation disorder, clearly associated with muscarinic receptor antagonism but not with GABA. When we tested zolpidem for its effects at a large range of targets, we found that in addition to engagement to the GABA-A receptor (AC50 = 46 nM) it also bound to the M1 muscarinic receptor with high potency (AC50 = 210 nM). Eletriptan is a highly selective HTR1A receptor agonist for the treatment of migraine^47^. To our knowledge, there is no evidence that it has pharmacodynamically relevant muscarinergic engagement which could explain the predicted accommodation disorder in association with the M2 muscarinic receptor.

### Case study: Citalopram off-target-ADR associations

Citalopram is a SSRI antidepressant used to treat anxiety disorders and other psychiatric conditions^48^. While catecholamine uptake was broadly investigated with SSRIs^49^, there is little knowledge about their engagement with monoaminergic and other CNS receptors, in contrast to other antidepressants.^50^ We found several ADRs that could be associated with dopaminergic engagement: movement or extrapyramidal disorders, psychotropic disorders and endocrine adverse reactions (Table S7). Our results indicate modest activity at D1 and D3 (AC50 = 4 µM and 5 µM, respectively), but not D2 receptors (AC50 > 10 µM). The major metabolite of citalopram, desmethyl-citalopram, has been reported to have similar binding activity at the D3 receptor^51^.

There is published evidence that SSRIs, including citalopram, are rarely associated with TD. The present hypothesis is that this is related to an indirect anti-dopaminergic effect induced by increased levels of serotonin^52,53^. However, there is indirect support for interaction of citalopram with the dopaminergic system. Citalopram induces the upregulation of dopamine D1, D2 and D3 receptor mRNA levels in the rat nucleus accumbens^54^ which is an essential part of the rewarding and primary motivational CNS network. The results indicate that alterations in the availability of neurotransmitters at synapses induced by citalopram are strong enough to induce immediate and long-lasting adaptive changes in the neuronal network. However, the strongest effect was observed with the D2 receptor^55^. This finding also raises the possibility, that homo- and/or hetero-dimerization^56^ of the dopamine receptors might occur due to citalopram treatment and potentially results in D2 upregulation^57^ suggesting that antidepressants can induce adaptive changes in the brain.

Based on the engagement with the dopaminergic system, it is not surprising that citalopram also has psychotropic side effects similar to that of atypical antipsychotic drugs. However, this class of ADRs is difficult to differentiate from symptoms associated with the treated disease^12^. Also, the serum concentration of citalopram is affected by concomitant treatment with neuroleptics, benzodiazepines, and tricyclic antidepressants and by the age of the patients^58^. These conditions have two important effects on the ADR association of citalopram, namely the concomitant treatment could cause the side effects and the elevated concentration of citalopram could bring its weak effects at the serotonin and dopamine receptors in coverage. Thus, careful analysis is needed to link common ADRs of psychotropic drugs to citalopram.

Finally, we have encountered information on hormonal changes, namely hyperprolactinemia, an endocrine disorder that is associated with risperidone, an atypical antipsychotic drug^59^. It manifests in galactorrhea and gynecomastia particularly prominent in boys treated for irritability associated with autism^60^. In a previous study, we linked high volume of hyperprolactinemia/gynecomastia reporting in FAERS to strong engagement of risperidone to serotonin and dopamine receptors, transporters with a narrow safety window^12^. However, endocrinal ADRs are rarely associated with antidepressant drugs (SSRIs) and monoamine oxidase inhibitors (MAO-I)^61^. The HRH2 activity noted herein may therefore contribute to ADR of hyperprolactinemia reported in SIDER.

## DISCUSSION

*In vitro* safety pharmacology assays are an important tool in lead optimization and risk assessment prior to human studies. With the goal of interpreting results for new chemicals in context, we created the Secondary Pharmacology Database (SPD) over a multi-year period to characterize approximately 2000 drugs. In this work, we present a comprehensive analysis of bioactivity vs. ADR relationships using uniform and standardized assay protocols. Many of these assays are offered by commercial vendors, allowing application of the SPD for probing safety characteristics of newly synthesized compounds.

By comparing results from our database to freely available (ChEMBL, DrugCentral) and subscription resources, we found that ca. 95% of assay results were unique to the SPD. This proportion was highest among inactive results but remained ca. 36% for results with AC50 < 1 µM. When activity results in SPD were also described in the public resources from literature curation, literature-reported AC50 values tended to be smaller (i.e., more potent). These observations are consistent with a bias towards publishing active results.

The availability of inactive results (typically AC50 ≥ 30 µM) for many drug vs. target pairs allowed us to test the association between presence or absence of drug ADRs (either text-mined from labels – the SIDER database – or FAERS) and the presence or absence of activity at a given target. When evaluating the statistical significance of literature-reported target vs. ADR relationships, we found 64% with no support. This varied significantly by target, suggesting that smaller panels of assays based on carefully selected targets could be used at earlier stages of lead optimization. Our modelling suggested that many associations lacking support reflect limitations in the dataset (i.e., too few potent drugs for the target). However, 128 associations flagged by the model as unlikely to be significant have strong support from our data (p < 0.001). This suggests that many of the literature associations lacking support may have modest predictive utility (low effect size), even if substantiated in larger datasets. Contrary to our expectations, we found more relationships to be supported by unadjusted AC50 values vs. human Cmax-derived safety margins. From a practical perspective, this facilitates risk assessment early in lead optimization when human Cmax estimates may not be available.

We systemically analyzed target vs. ADR pairs using our database, identifying potentially novel associations. Statistical analyses linking targets to ADRs are highly confounded by polypharmacology, whereby ADR risk might be attributed to target activity correlated with the causal risk drivers. Prior investigations using publicly available assay data either did not control^15^ or clustered similar associations to identify putative drivers^10^. We attempted to control known associations by performing penalized multivariate selection using all individually significant assays, including lower significance associations reported in the literature. This approach eliminated ca. 800 associations, increasing the likelihood that those retained are causal.

This approach has several limitations. ADRs from SIDER and disproportionality analyses on FAERS data may be confounded by drug indication^12^. Further, since many druggable proteins were not included on our panels, the absence of the causal proteins would fail to deselect non-causal proteins. For instance, 12 associations involving the protein kinase KDR are listed as “potentially novel” in Table S6. These include “stomatitis”, “gastrointestinal perforation” and “malignant neoplasm progression”. These associations may represent general kinase mediated ADRs, with the latter illustrating confounding of ADRs due to drug treatment vs. symptoms associated with the treated disease.

An illustrative example concerns associations between targets and “electrocardiogram QT prolonged”. Logistic regression was applied to several assays, which are either described as risk factors for long QT in the literature reviews (ADRB2, HRH1, KCNH2 and SLC6A2) or were identified by univariate analyses (ADRA1A, DRD2, DRD4, HRH2 and HTR2B). Among the literature-reported associations, activity at the hERG channel (KCNH2) is thought to be the primary risk factor. The penalized modelling approach retained ADRA1A, DRD2, HRH1 and KCNH2 as non-redundant variables. The role of all but KCNH2 is controversial. Several drugs devoid of activity at KCNH2 (i.e., AC50 > 10 µM) are annotated with long QT in SIDER or FAERS and support these associations (i.e., AC50 < 1uM): ADRA1A (aialfuzosin, clonidine, mianserin, olanzapine, quetiapine), DRD2 (amisulpridem, olanzapine, quetiapine) and/or HRH1 binding (cetirizine, mianserin, mirtazapine, olanzapine, quetiapine, valproic acid).

There is overwhelming evidence of cardiac arrhythmias caused by H1 histamine receptor antagonists. While broad range of antihistamines could cause QT prolongation either on their own or in combination with other drugs, there has been an emerging common denominator, which makes this class of medications prevalent to cause cardiac arrhythmias. That is hERG channel inhibition, which has been reviewed extensively during the past two decades^62^. All H1 antihistamines and/or their metabolites – with very few exceptions - have direct hERG effect with various potency. Their effect might be exacerbated by high exposure because of co-administration of drugs interfering with the metabolism of the antihistamines. Alternatively, combination therapy with other hERG inhibiting drugs could synergize their effects. In summary, caution is needed when ubiquitous off-target effects appear in a class of drugs aiming at the same therapeutic target.

Throughout this work, we labelled as “novel off-target” any activities in SPD not reported in ChEMBL, DrugCentral or the subscription resources containing curated pharmacological activity results. Curation-based resources have limited journal coverage, and results we claim as “novel” can sometimes be found with manual searches (e.g., PubMed, Google Scholar). For example, lurasidone and vandetanib are annotated with the ADR “Torsade de pointes” and were found to have hERG AC50 values of 0.53 and 0.35 µM, respectively. These activities are not reported in the sources we considered, but are reported in the literature^63,64^. A condition for practical large-scale analyses of pharmacological activity results is inclusion in commonly used databases. As such, activity results from SPD are a significant addition to existing resources summarizing bioactivity of drugs.

## METHODS

### In vitro safety pharmacology assays

Compounds were obtained from the Novartis Institutes of Biomedical Research (NIBR) compound library and tested in a panel of in vitro biochemical and cell-based assays at Eurofins and/or NIBR in concentration-response (8 concentrations, half-log dilutions starting at 10 or 30 µM). Assay formats varied from radioligand binding, to isolated protein, and cellular assays. Normalized concentration response curves were fitted using a four-parameter logistic equation performed using software developed internally (Helios). The equation used is for a one site sigmoidal dose response curve: Y=A+((B-A)/(1+((X/C)^D))), where A=min, B=max, C=IC_50_, D=slope. By default, min is fixed at 0, whereas max is not fixed.

When drugs had no significant biological activity at the highest concentration tested, the AC50 was reported with qualifier “>”; for example, an AC50 is reported with qualifier “>” and AC50 value “30” when a compound exhibits no significant activity at concentrations up to 30 µM. Where curve fitting produces an AC50 value below the highest concentration tested, activity is reported with qualifier “=”.

### Mapping chemicals to DrugCentral structure IDs

Multiple chemical substances were tested in the SPD assays. Different substances include distinct lots sourced from chemical vendors or synthesized internally of a given drug, different salt forms, and/or metabolites of the parent drug. In preparation of the dataset released with this study, SMILES representation of substances from the NIBR chemical database were de-salted and converted to InChI keys using RDKit’s MolToInchi function (https://rdkit.org/; accessed 09/22/2021).

The DrugCentral PostgreSQL database dated 09/18/2020 was downloaded (https://drugcentral.org/download; accessed 09/22/2021). The InChI key consists of three parts separated by hyphens, of 14, 10 and 1 character(s), respectively. These correspond to the connectivity information (or graph; 14 characters), remaining layers (10 characters) and protonation state (1 character). For each NIBR structure, matches to DrugCentral were attempted at multiple levels of decreasing stringency: 1) perfect matches: the InChI key obtained on the DrugCentral SMILES matches the key from a Novartis SMILES, 2) match without the protonation part of the key, 3) match using only the graph part of the key, but require a name or synonym match, and finally 4) try to match on name or synonym, and review structures. Level 4 matches are common with complex drugs, such as natural products, where drawing errors occurred during registration of substances. Matches involving names and/or synonyms compare those from DrugCentral to names assigned as part of the NIBR substance registration. Note that DrugCentral sometimes includes both parent drugs and metabolites, both of which were used in the matching process.

### Dataset summarization

Substances sharing the same InChI key tested in multiple assay runs were summarized into a single numeric AC50 for a given InChI vs. assay pair. When one or more AC50 values had qualifier “=“, the geometric mean was computed and reported with qualifier “=”; N summarized indicates the number of averaged AC50 values, and N total indicates the total number of AC50 values for the InChI vs. assay pair, including AC50 values with qualifier “>” excluded from the geometric mean computation. In the absence of any AC50 value with qualifier “=”, the largest value among those with qualifier “>” was retained. For instance, the AC50 values of “>1 µM” and “>30 µM” are summarized as follows: qualifier “>”, numeric AC50 value “30”, N summarized “2” and N total “2”.

### Defining assay groups

The SPD database was created over several years, with some targets having multiple assay protocols employed. Each protocol was assigned a unique identifier. Changes in protocol often result in the creation of a “new” assay, designated with a new identifier, even when both assays effectively measure the same biochemical event. Examples include changes in radioligand, measurement technology (e.g. filter binding vs. TR-FRET), outsourcing to a contract research organization (CRO), etc. To maximize power for detecting statistically significant relationships and simplify analysis, we defined assay groups that combine assays resulting in concordant AC50 values when comparing results on the same drug substances.

Concordance analysis was performed by evaluating agreement of assay results where at least 10 compounds were tested in each pair of assays having the same target and mode (e.g., both binding, agonist or antagonist assays). Qualitative agreement (<10 µM, ≥ 10 µM) was assessed by calculating the sensitivity of each assay for detecting actives from the other, with a minimum of 0.5 for both assays (i.e., # active in both assays / # active in assay1 ≥ 0.5 and # active in both assays / # active in assay 2 ≥ 0.5, and Pearson R ≥ 0.7 calculated on 10 or more log AC50 values, where both results had qualifier “=”. Assay pairs with insufficient overlapping test compounds were not merged. Viewing assays as nodes and concordant assay pairs as edges, all nodes within a connected graph were merged, even if some assay pairs within the group fell below our concordance cutoff (or lacked sufficient overlapping pairs to assess). To increase the number of overlapping test compounds for a pair of assays, this analysis was performed on all results available for the assays, including proprietary compounds not included in the supplement. Dataset S1 is provided at the level of both individual and grouped assays.

Within each assay group, the assay supplying the largest proportion of results was designated as the preferred assay. When results were available for the preferred vs. other group assays, results from preferred assay were used in downstream analyses.

### Integration of DrugCentral, ChEMBL and subscription activity results

Activity data from DrugCentral, including annotation of targets as drugs’ mechanism of action (MOA) were obtained from the “act_data_full” table for human and mammals (rat, mouse, cow, guinea pig, rabbit, pig, sheep, dog, chicken, and monkey). Drugs from the SPD were mapped using the DrugCentral structure ID as described above. For each DrugCentral target ID, the Swissprot identifier from the “target_component” was mapped to Entrez gene IDs using the gene2accession file from Entrez gene, or the Uniprot ID mapping tool (https://www.uniprot.org/uploadlists/; accessed 09/23/2021) in the absence of an Entrez match. Finally, any non-human Entrez gene IDs were mapped to the human ortholog using a compilation of associations from Ensembl, Homologene, RGD, and MGI. For SPD, assays using non-human proteins were represented with the human Entrez gene ID. Each drug vs. assay pair from SPD was annotated with the median AC50 for all DrugCentral activity records with the same structure ID and human gene ID. This matching did not consider activity mode (inhibitor, antagonist, etc. - action_type in “act_data_full”), because it was undefined in most cases.

When defining on-target activity, the DrugCentral “act_type” variable defining a drug’s mode of action at the target was compared to the assay mode. For functional assays, only drugs annotated as having the same mode as the assay were retained (e.g., agonist drugs for agonist assays). For GPCR and nuclear receptor assays having “binding” or “inhibition” modes, agonist drugs were removed. Antagonist drugs were retained because binding and functional antagonist readouts are correlated on our panels.

ChEMBL version 27 was downloaded as a PostgreSQL database. To increase the number of matches while allowing variation in structural representations (e.g., ignoring chirality, structure drawing errors, etc.), ChEMBL compound identifiers provided by DrugCentral were supplemented with those matching the SPD InChi keys identified using the multi-criteria match described above. ChEMBL targets identified with Uniprot accession numbers were mapped to SPD assays as described above. Activity types and units suggestive of multiple concentration testing were converted to AC50 values in µM units (pM, nM, µM, mM, M); results with units of ng/mL, pg/mL, ug/mL were converted to molarities using the drug free base RDKit molecular weight. ChEMBL results suggestive of single concentration testing (units of %) were considered separately. Multiple ChEMBL drug vs. target values were summarized as the median, separately for AC50 and single concentration results.

Clarivate Cortellis (https://clarivate.com/cortellis/solutions/pre-clinical-intelligence-analytics/; accessed 8/3/2021) and Excelra GOSTAR (https://gostardb.com/gostar/; accessed 8/3/2021) are subscription-based resources similar to ChEMBL and DrugCentral. The same processing applied to ChEMBL was used to identify median AC50 values for SPD drug vs. assay pairs.

### Selecting a representative SPD result for each drug vs. assay group pair

Multiple InChi keys are available for certain drugs, and multiple assays within assay groups. To simplify analysis, we denoted one result as representative among all available for a given prescribed (parent) drug vs. assay group pair. For a given drug vs. assay group pair, the algorithm selects assay results for the active metabolite over any collected for the parent drug (e.g., take activity results for enalaprilat when describing activity of enalapril). The algorithm favors highly quality structural matches between DrugCentral and SPD and selects from the preferred assay within the assay group. Detailed logic is available in the Jupyter notebook make_all_prescribable_drug_activity_dataset.ipynb. The selected activity records have column “representative_result_drug_assay_group” = TRUE in Dataset S1.

### Human drug exposure (Cmax) and plasma protein binding

The maximal drug exposure (Cmax) at highest approved dosage was curated from the primary literature for 487 drugs. To broaden the dataset, we extracted human Cmax values from Pharmapendium (https://pharmapendium.com; accessed 10/26/2021). These are often from heterogeneous sources, and include results from lower doses, metabolites, pediatric studies, Cmax at steady state, etc. Selecting a representative value could be achieved by calculating the median, average, 3^rd^ quartile or 90^th^ percentile. We employed the 487 manually curated results as a reference set and found maximal correlation when Pharmapendium values were summarized as the 3^rd^ quartile. This supplied a further 451 drug Cmax. Finally, we used values reported in Smit et al^15^ for a further 146 drugs, yielding a dataset of 1084 marketed drug Cmax values (Table S1).

To calculate free Cmax values, we employed a similar approach for compiling plasma protein binding (PPB) %: primary literature curation (572 drugs), Pharmapendium (332), DrugCentral (90) and Smit et al (52), providing a dataset of 1045 plasma protein binding. For Pharmapendium, results reported as albumin or glycoprotein binding were excluded because correlation vs. our curated dataset was low; the 3^rd^ quartile provided the highest concordance vs. curation, as observed for Cmax.

Cmax(free) was determined as the product (100-PPB%) x Cmax(tot) for 940 drugs having both parameters available.

### Adverse drug reaction annotation using FAERS and SIDER

Annotation of drugs as being positive or negative for MedDRA-coded adverse drug reactions was performed using two sources. In the absence of freely available high quality manual curation of adverse drug reactions and their frequency from labels for all FDA approved drugs^65^, surrogate approaches must be employed. The FDA adverse drug reaction reporting system (FAERS) is often used in pharmacovigilance research to detect adverse drug reactions. In this work, we used FAERS data from DrugCentral^24^ without any further post-processing. Adverse drug reactions with log likelihood ratio (LLR) ≥ 5 times the drug-specific threshold value (llr_threshold) were deemed positive for a given drug^14^, otherwise the drug was labelled as negative for the adverse drug reaction.

The SIDER database provides annotation of drug ADRs obtained from text mining applied to drug labels^16^. SIDER uses drug annotation from STITCH, a sister database (http://stitch.embl.de/download/chemicals.v5.0.tsv.gz; accessed 09/23/2021). SMILES from STICH were converted to InChi keys and matched to DrugCentral structures using the multi-criteria matched described above. The mapping is provided in Table S9.

Drug ADRs were obtained from the file meddra_all_se.tsv.gz available from the SIDER website (http://sideeffects.embl.de; accessed 09/24/2021). Drugs were mapped to DrugCentral using the stereo_id using Table S9. SIDER ADRs are labelled using UMLS CUIs; they were mapped to MedDRA preferred term (PT) codes using the UMLS REST API (https://documentation.uts.nlm.nih.gov/rest/home.html; accessed 09/24/2021). A version of meddra_all_se.tsv, using DrugCentral struct_id and MedDRA PT codes is provided as a supplementary dataset with the Jupyter notebooks (final_sider_map_to_drugcentral_meddra.txt).

### Creating adverse drug reaction training sets

For each ADR, positive drugs (causing the ADR) and negative drugs (not causing the ADR) must be defined. To study the association between ADRs and assays, ADR terms from SIDER and FAERS reported as MedDRA preferred terms (PTs) were mapped to MedDRA high level terms (HT) and high-level group terms (HG). The mappings were obtained using the UMLS API. Mapping to higher terms results in more drugs labelled as positive (i.e., higher power to detect an effect), but potentially combining PTs with distinct target (assay) risk factors. We therefore modelled relationships at 3 levels: PT, HT and HG.

Depending on the level of MedDRA terms, different strategies were employed for defining ADR negative drugs (i.e., drugs that do not cause a given ADR). For HGs, any drug not positive for one of the underlying PT terms was considered a negative. For HTs, any drug not positive for the term *and* not positive for a sister HT under the current HG was considered a negative. For PTs, any drug not positive for the term *and* not positive for a sister PT under the current HT was considered negative. This strategy was used to avoid labelling as negative a drug that produces a similar adverse drug reaction to the one under study. For example, negatives for PT 0044066 (Torsade de pointes) would exclude drugs that are positive for PT 10047302 (Ventricular tachycardia), because both PTs share the HT 10047283 (Ventricular arrhythmias and cardiac arrest). The Jupyter notebook make_ADR_training_sets.ipynb was used to automate the definition of positive and negative drugs for each PT, HT and HG term.

Because drugs sharing an active metabolite may have different ADR annotations (e.g., betamethasone dipropionate vs. betamethasone valerate), each was treated separately. Annotation of drugs with a selected activity record from SPD for each assay was performed using the multi-criteria approach described above, using the Jupyter notebook make_ADR_vs_activity_dataset.ipynb. It should be noted that this differs from make_all_prescribable_drug_activity_dataset.ipynb, where we selected a single representative drug form (betamethasone) among all those tested.

### Univariate adverse drug reaction vs. assay association

To establish the strength of association between drugs’ status for adverse drug reactions (Boolean) and assay activity (continuous measures of AC50, free and total margin), the Kruskal-Wallis test and ROC AUC computation was performed for each ADR vs. assay pair.

Activity measures of AC50, total margin and free margin frequently have qualifier ‘>’, indicating that measured AC50 was estimated to exceed the highest concentration tested in the assay. The maximum tested concentration of 10 µM and 30 µM were employed for most assays. To calculate a rank-based association test between assays and ADRs, it was necessary to select an AC50 cutoff and replace all values in excess with the cutoff value (truncating). Values with qualifier ‘>’ but AC50 below the cutoff were excluded. For AC50 values, the numeric distribution for qualifier ‘=’ and ‘>’ were largely non-overlapping, the natural cutoff is 10 µM or 30 µM depending on the assay, and few values needed to be truncated or excluded. Because drug total and free Cmax vary over a wide range, safety margin distributions overlap significantly for qualifier ‘=’ and ‘>’. This makes the selection of cutoff more difficult: too low and one loses the ability to distinguish ranks for drugs with safety margin above the cutoff, but too high and one must exclude from analysis many values with qualifier ‘>’ below the threshold (and hence a loss of power). We performed tests using cutoffs of 10 µM and 30 µM for AC50, 2 and 10 for total margin, and 10 and 100 for free margin.

For assessing the statistical significance of literature-reported assay vs. ADR associations (see below; Table S10), we retained the threshold giving the smallest (most significant) KW p-value. For systematic analysis of all assay vs. ADR pairs, we selected a single threshold per assay in order to minimize the number of tests performed (i.e., increasing the false discovery rate). For each assay and activity measure (AC50, total margin, free margin), the cutoff providing the largest total number of assay vs. ADR associations with ROC AUC ≥ 0.7 and KW p-value ≤ 1e-06 was selected and used for all further analyses. The higher cutoffs (30/10/100 for AC50/total margin/free margin) were generally selected (37 vs. 23 assays for AC50, 28 vs. 14 assays for total margin and 35 vs. 5 assays for free margin cutoff).

Excluded from analysis were all combinations of assays, activity measures (AC50, free margin, total margin) and ADRs not meeting the following criteria: 10 or more positive drugs (i.e., drugs with the ADR), 50 or more negative drugs and 10 or more non-qualified activity values. Univariate analyses were performed using the Jupyter notebook calc_ADR_vs_assay_score.ipynb.

### Multivariate modelling of adverse drug reactions

Multiple assays and activity measures (AC50, total margin, free margin) may show significant association with a given ADR. Because assay activity is often correlated across related targets (e.g., targets with similar binding pockets), variable selection strategies can be used to select a smaller number of assay and activity measures among all those which met our univariate threshold (KW p-value ≤1e-06 and ROC AUC ≥ 0.7). We used L1-penalized (Lasso) logistic regression to model each ADR outcome (positive or negative) using the subset of assay + activity measures selected by univariate analysis. AC50 and margins were log10 transformed prior to modelling.

When modelling ADR outcomes with multiple assays, missing activity values occur when some drugs were not tested in all assays. For each ADR, we required individual assays to reach 70% or greater coverage compared to the assay with maximal drug count; missing values were imputed by using the median.

For each dataset (ADR class as dependent variable and assay activities as independent variables), a sequence of penalties ‘c’ was generated. Average and standard error of ROC AUC at each penalty were determined using 50 trials of leave 20%-out cross-validation. The smallest ‘c’ (or largest penalty) producing a model with ROC AUC within 1 standard error of the best model was selected. This led to the creation of models with few variables, in order to discern the principal contributors to ADR risk. Variables with zero coefficient have no significant role in explaining odds of being positive for a given ADR, after accounting for the contributions of variables with non-zero coefficients.

Because small values of AC50 or margins indicate activity of drugs in an assay, variables with large negative coefficients in the logistic regression model represent assays for which increasing activity results in higher odds of being positive for a given ADR. For coefficients ≥ −0.08, the variable was tagged as not in the model. This corresponds to interpreting a 10-fold decrease in AC50 or margin being associated with a smaller than 20% increase odds ratio of observing the ADR. There were occurrences of coefficients > 0.1, i.e., indicating decreased risk of the ADR for activity in the assay. These were almost exclusively in models where an expected negative association was present for another activity parameter of the same assay (i.e., AC50 had a large negative coefficient and free margin had a small positive coefficient). These were considered excluded from the model (i.e., coefficient of 0 in Table S6). Multivariate analyses were performed with the Jupyter notebook build_ADR_vs_assay_model.ipynb.

### Literature-derived target vs. ADR associations

Target-ADR relationships, as published in 3 key reviews, were obtained from the supplementary material in Smit et al^15^. Because they did not provide the direction of association (target activation vs. inhibition), we reviewed associations from the 3 publications. Smit et al. mapped terminology from the reviews to MedDRA preferred terms (PT). In reviewing their results, we added some missing associations and refined mappings to MedDRA codes. These are denoted as “pre-analysis” supplemental terms in Table S10.

Because selection of MedDRA PTs from the literature reviews may differ from their representation in SIDER or FAERS, we examined the frequency of each literature PT code in SIDER and FAERS. Some terms with suspiciously low frequency triggered searches for better terminology. For instance, ‘Intestinal transit time decreased’ is a valid MedDRA PT used in Lynch et al, however it is not used in SIDER or FAERS. However, both ‘Gastrointestinal disorder’ and ‘Diarrhea’ were identified as substitutes. These additional mappings were added to the Literature vs. MedDRA PT term mapping.

For each combination of MedDRA code and target from the literature, we examined the significance of the association in the SPD. Each association was tested across all SPD assays for the target (median 1, range 1-6), 2 sources (SIDER, FAERS), 3 activity measures (total margin, free margin, AC50) and 2 activity cutoffs for denoting active vs. inactive results (total margin 10 vs. 2, free margin 100 vs. 10, AC50 30 vs. 10 µM). As such, 12 to 72 tests (median of 12) were conducted per literature association, and we classified as “marginally significant” those associations with KW p-value between 0.05 and 0.001 (nominal p-value of 0.001 with 72 tests yields a Bonferroni-corrected p-value ~ 0.07). We only tested associations having at least 10 positive drugs and 50 negative drugs with available assay results; adverse drug reactions with counts below these thresholds were typically rare (e.g., death) and were classified as “not tested”.

Failure to achieve significance might be due to a poor selection of MedDRA PT for the ADR. For associations classified as marginal, not significant or not tested, we repeated the statistical testing described above for each PT that shares a given HT with the literature derived term. For example, Smit et al. mapped “urinary contraction” (given in the Bowes review as an ADR for CHRM3 activation) to the MedDRA PT “Bladder spasm” (MedDRA 10048994). Our dataset only contained 6 drugs annotated with this ADR (classification “not tested”). However, several MedDRA PTs sharing the same HT met our criteria of KW p-value ≤ 0.001 and ROC AUC ≥ 0.6. Ordered by increasing p-value (most significant first), these include “Urinary retention” (10046555), “Urinary hesitation” (10046542), “Micturition disorder” (10027561), “Strangury” (10042170). The selection algorithm sorted all related terms by p-value, accumulating the number of tests performed, and stopping at the first satisfying the above criteria. In Table S5, the association of CHRM3 activation with “urinary contraction” (literature ID 364) contains the p-value and ROC AUC for “Urinary retention” and is classified as highly significant (p≤1e-06), with distance 1 (i.e., the significant PT and starting PT are connected via 1 intermediate in the network, via the shared HT).

When no PT terms sharing a given HT were identified, terms sharing a high-level group term (HG) were examined. These are encoded as distance 2 in Table S5 (i.e., the starting and significant term are linked via two intermediates: HT then HG. For both the HT and HG expansion, we used a two-step process: first identifying possible related terms, then manually reviewing and confirming them. This is especially important for distance 2 relations where very broad (and sometimes opposite) effects are grouped at the HG level. Only 14 target ADR pairs were found significant via a shared HG term (distance = 2), but not distance 0 or 1; 11 of these used the term “Ileus paralytic” (10021333) ascribed to “constipation”, “gastrointestinal motility decreased”, “gastrointestinal transit decreased”) reported in the 3 reviews.

The Jupyter notebook calc_lit_AE_vs_assay_score.ipynb was used to perform this analysis.

The final set of literature-derived target vs. ADR annotations, including those obtained from Smit et al. and our additions via the common HT and HG terms, are provided in Table S5. Tabulation as “significant”, “marginally significant”, “not significant” or “not tested” in results uses the strongest association for any of the individual MedDRA PTs regardless of their source. Developmental ADRs from Lynch et al^9^ are included in Table S5 for completeness but were excluded from result summaries described throughout. Because these are rare effects, only 3 of these associations meet the criteria of 10 positives and 50 negatives in SIDER and/or FAERS.

### Logistic regression modelling of variables associated with significance of literature associations

Establishing statistical significance of a given target vs. ADR pair requires having sufficient number of drugs that are positive and negative for the ADR, and a sufficient number of drugs with measurable activity in the assay. We selected minimal but arbitrary requirements of 10 positives, 50 negatives and 10 or more non-qualified activity values to test the association. Failure to find significance may simply reflect the limited power of the dataset, i.e., the above cutoffs were not set high enough.

Because the majority of significant literature-reported associations were supported by the AC50 activity measure (rather than free or total margin), we repeated the selection process described above to identify the most significant MedDRA code, source (SIDER or FAERS), assay and activity cutoff for the AC50 activity measure only. This avoided combining AC50 and margin-based activity measures having different scales, and for which standard percentile values would be non-comparable (free margin = 1 and AC50 = 1 µM are not comparable).

Literature-reported target vs. ADR pairs were classified as “significant” (222 pairs) or “non-significant” (497 pairs), using the criteria KW p-value ≤ 0.001 and ROC AUC ≥ 0.6. Since this analysis used only the AC50 activity measure, there are fewer “significant” associations compared to Table S5 (which used AC50, free and total margin). To identify families of ADRs more or less likely to be significant, MedDRA PTs were mapped to system organ classes (SOCs), and each literature association was annotated as assigned to (1) or not (0) a given SOC. Some PTs map to multiple SOCs (e.g., “Metabolism and nutrition.disorders”, “Endocrine disorders”). Several summary statistics that capture the proportion of drugs with potent activity in the assay were selected: percentile values (2.5, 5, 10, 25 or 50^th^; e.g., assays with many potent drug activities will be represented with smaller percentile values), count of AC50 values less than or equal to 100 nM, 500 nM or 1uM, and count of drugs with assay results that are positive or negative for an ADR. The dataset used for this modelling is provided in Table S11.

Lasso-penalized logistic regression modelling was performed using the R package Glmnet using default parameters [glmnet(x,y,family=“binomial”)] and the AUC metric for cross-validation [cv.glmnet(x,y,family=“binomial”, type.measure = “auc”)].

### Novel assay results vs. known drug adverse drug reactions

Novel SPD activity results, i.e. drug-target pairs not reported in the sources we considered, were compared to known drug ADRs to determine if these novel activities might explain the ADRs. The following criteria were applied: 1) drug has AC50 < 5 µM at a given target that is not reported in ChEMBL, Drug Central or the subscription resources, 2) drug is known to cause a given ADR according to SIDER and/or FAERS, 3) literature reported target vs. ADR relationship is significant in SPD (p ≤ 0.001) on one of more of AC50, free or total margin, 4) the drug’s activity on one or more of those significant measures is in the top 3 quartiles among all drugs active in the assay and having the ADR, 5) the drug’s on-target activity is not associated with this ADR, 6) any known off-target activities associated with the ADR have significantly lower potency compared to the new activity (≥ 10 fold). The final criteria avoid flagging novel weak activities unlikely to make significant additive contributions to the ADR risk.

## List of Supplementary Materials

### Figures

S1. Statistical significance of literature-reported target vs. ADR associations by activity measure and source.

S2. Identification of attributes associated with significance of literature-reported target vs. ADR pairs.

S3. Distribution of significant literature-reported target-ADR pairs.

S4. Comparison of literature-reported target-ADR pairs assessed on free margin vs. AC50.

### Tables

Provided in Excel format, SI_tables.xlsx

S1. Unique drug substances tested in safety pharmacology assays

S2. Safety pharmacology assays used for characterizing marketed drugs S3. Safety margin characteristics of on-target drug interactions

S4. Comparison of on-target indications vs. top-10 indications associated with the novel off-target activity

S5. Significance analysis of literature reported target vs. adverse event relationships

S6. Assay vs. AE pairs from univariate analysis and annotation with multivariate model coefficients

S7. Details of NSPD activities not reported in literature sources potentially associated with drugs’ known clinical adverse events

S8. Summary of NSPD activities not reported in literature sources potentially associated with drugs’ known clinical adverse events

S9. Mapping of chemical identifiers from SIDER / STITCH to DrugCentral

S10. Literature-reported target vs. AE associations from 3 reviews and annotation with MedDRA terminology

S11. Dataset of most significant AC50 result for each literature-reported target vs. ADR relationship

### Code

www.github.com/Novartis/SPD

Jupyter notebooks described in methods:

make_all_prescribable_drug_activity_dataset.ipynb

make_AE_training_sets.ipynb

make_AE_vs_activity_dataset.ipynb

calc_AE_vs_assay_score.ipynb

build_AE_vs_assay_model.ipynb

calc_lit_AE_vs_assay_score.ipynb

### Data

www.doi.org/10.5281/zenodo.7378746

Dataset_S1.xlsx: SPD activity database

Files used within Jupyter notebooks:

AE_vs_assay_pairs_logistic_modelling_update.txt

assay_group_vs_gene_map.txt

assay_vs_param_thresholds_update.txt

codes_vs_names.txt

combined_faers_sider.txt

drugs_for_lit_targets.txt

final_sider_map_to_drugcentral_meddra.txt

final_summarized_activity_data_pub.txt

lit_target_AE_pairs.txt

meddra_pair_synonyms_reviewed.txt

merging_assays_preferred_annotation_pub.txt

parent_to_metabolite_map.txt

terms_vs_parents.txt

## Supporting information

Secondary Pharmacology Database

## Acknowledgments

We thank Jonathan Moggs and Greg Friedrichs for their support. We also thank the Translational Medicine Data Science Academy team at NIBR.

## Author contributions

A. F., L.U. provided the secondary pharmacology material, J.S., D.Y. and A.F. analyzed and interpreted the data. J.S. and L.U. drafted the manuscript. All authors critically reviewed the manuscript.

## Data and materials availability

The data associated with this publication is freely available at the Zenodo repository (www.doi.org/10.5281/zenodo.7378746). In addition, the associated Python code in form of six separate Jupyter Notebooks described in the Materials and Methods section of the manuscript is freely available on GitHub (www.github.com/Novartis/SPD).

