## Supplementary Materials and Figures for "A novel preclinical secondary pharmacology resource illuminates target-adverse drug reaction associations of marketed drugs"

**Supplementary Results**

**Supplementary Methods**

**Supplementary Figures**

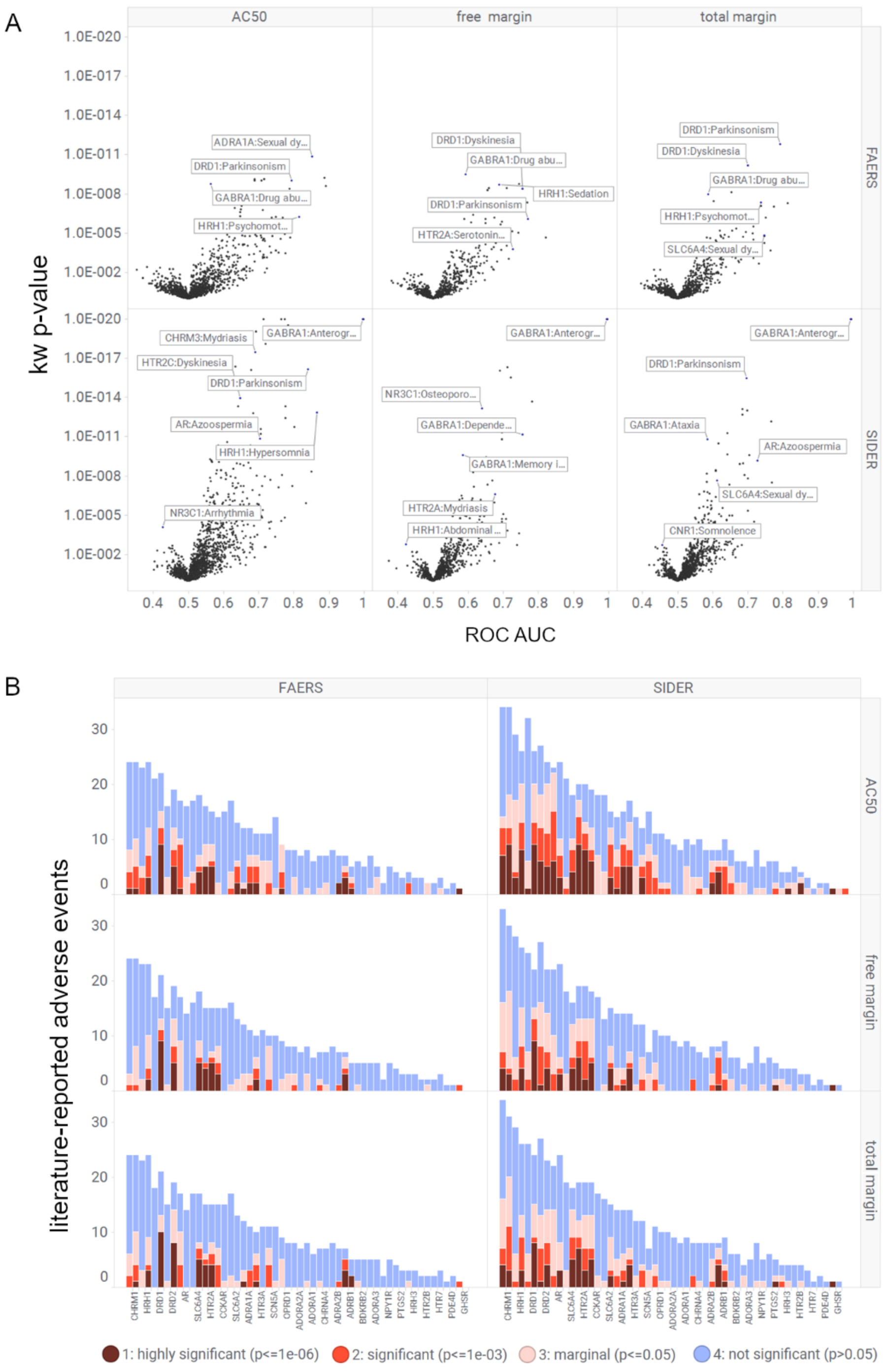


**Fig. S1. Statistical significance of literature-reported target vs. ADR associations by activity measure and source.** A) Comparison of KW p-value vs. ROC AUC by activity measure (AC50, free margin or total margin) vs. source (SIDER, FAERS) for literature associations. B) Total number of adverse drug reactions reported across targets, distinguished by level of statistical significance observed.

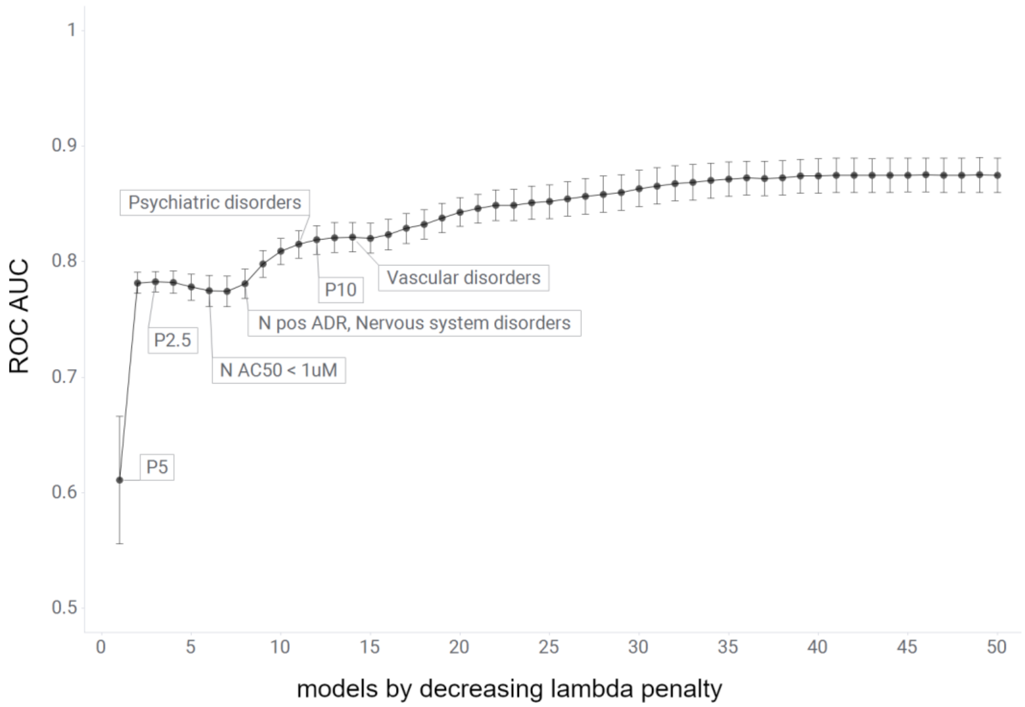


**Fig. S2. Identification of attributes associated with significance of literature-reported target vs. ADR pairs.** Literature target-ADR pairs were labelled as "significant" ($p < 0.001$ and ROC AUC $\geq 0.6$) or "non-significant" (all others) based on the KW-test and ROC AUC analysis. Penalized (lasso) logistic regression was used to classify outcomes, with models consisting of 5 variables having cross-validated ROC AUC ~ 0.8. Variables selected for inclusion in the smaller models are labelled.

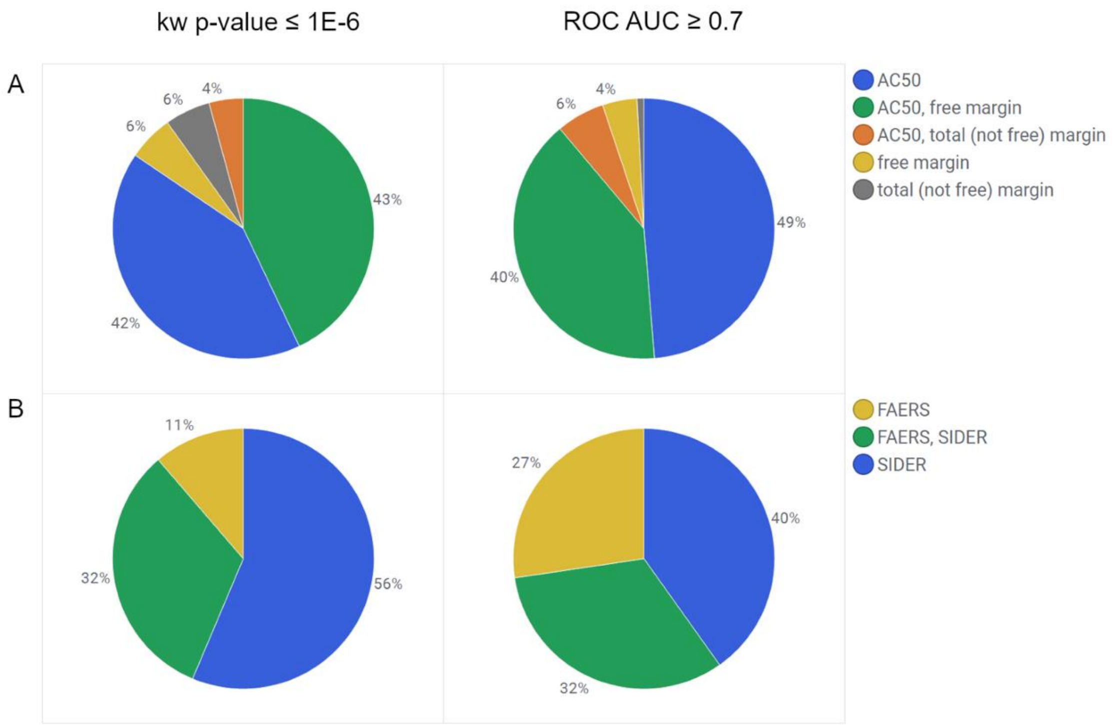


**Fig. S3. Distribution of significant literature-reported target-ADR pairs.** A) Distribution by activity measure: pairs significant for AC50 only (blue), AC50 and free margin (green – total margin not considered), AC50 and total margin (orange – i.e. not significant on free margin), free margin only (yellow - i.e. not significant on AC50, total margin not considered) and total margin only (gray). To simplify the number of categories, association on total margin was only considered when free margin was not significant. B) Distribution by ADR source: pairs significant in FAERS only (yellow), FAERS and SIDER (green), SIDER only (blue). Significance was assessed separately on KW p-value (left panel) vs. ROC AUC (right panel).

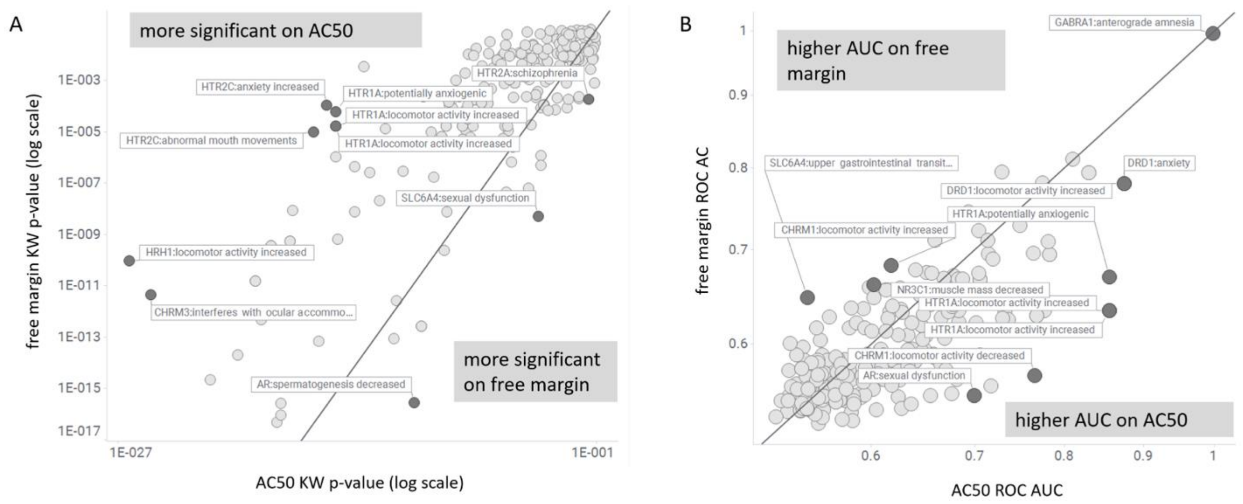


**Fig. S4. Comparison of literature-reported target-ADR pairs assessed on free margin vs. AC50**. A) comparison using KW p-value and B) ROC AUC. Selected target-ADR pairs are labelled as gene symbol: MedDRA name
